# Cell polarity opposes Jak-STAT mediated Escargot activation that drives intratumor heterogeneity in a *Drosophila* tumor model

**DOI:** 10.1101/2022.12.12.520127

**Authors:** Deeptiman Chatterjee, Fei Cong, Xian-Feng Wang, Caique Almeida Machado Costa, Yi-Chun Huang, Wu-Min Deng

## Abstract

In proliferating neoplasms, microenvironment-derived selective pressures promote tumor heterogeneity by imparting diverse capacities for growth, differentiation and invasion. However, what makes a tumor cell respond to signaling cues differently from a normal cell is not well understood. In the *Drosophila* ovarian follicle cells, apicobasal-polarity loss induces heterogenous epithelial multilayering. When exacerbated by oncogenic-Notch expression, this multilayer displays an increased consistency in the occurrence of morphologically distinguishable cells adjacent to the polar follicle cells. Polar cells release the Jak-STAT ligand Unpaired (Upd), in response to which, neighboring polarity-deficient cells exhibit a precursor-like transcriptomic state. Using single-cell transcriptomics, we discovered the ectopic activation of the Snail-family transcription factor Escargot (Esg) in these cells. We also characterized similar relationship between Upd and Esg during early follicular development, where the establishment of polarity determines follicle-cell differentiation. Overall, our results indicate that epithelial-cell polarity acts as a gatekeeper against microenvironmental selective pressures that drive heterogeneity.

## INTRODUCTION

Despite significant advances in anti-cancer therapeutics, poor prognosis of the disease is directly caused by the acquired resistance to therapy due to pre-existing Intra-Tumor Heterogeneity (ITH) and variations induced in response to the therapy. ITH is defined by the diverse morphological, genetic, epigenetic, transcriptomic and metabolomic states of neoplastic cells^1^. Currently, ITH is best understood from the perspective of genetic heterogeneity, where accumulating somatic mutations impart fitness advantage to the diverse cancer-cell lineages against selective pressures derived from the tumor microenvironment^2^. However, our knowledge of the non-genetic origins of ITH in spontaneous tumors remains somewhat rudimentary given the lack of suitable experimental models that reproduce ITH as observed in human tumors. This lapse is because the conventional animal models use powerful genetic combinations to drive reproducible tumorigenesis, causing the resultant tumors to rarely display naturally-occurring ITH as the cells represent the genotypes of only the winner-cell lineages. Therefore, better characterization of spontaneous ITH necessitates the use of model systems that represent the earliest steps of tumorigenesis.

In several tissues of *Drosophila*, tumorigenesis has been modeled by inducing the expression of oncogenic Ras or Notch along with the simultaneous loss of apicobasal-polarity genes *scribble* (*scrib*), *lethal (2) giant larvae (l(2)gl* or simply, *lgl*) and *discs large* (*dlg*)^3–6^, which is one of the earliest features of dysplastic tumors^7^. Polarity loss is also considered to be a founding event during Epithelial-Mesenchymal Transition (EMT) that grants plasticity to a cancer cell, thereby allowing it to exist in a multitude of metastable hybrid phenotypes^8,9^. While the tumors modeled in *Drosophila* do exhibit ITH^10^, complexities arising from rampant malignancy in these tissues obfuscate the study of individual cells. As an alternative, *Drosophila* ovaries provide a particularly favorable system to study ITH. Inducing polarity loss in the ovarian follicle cells promotes dysplastic multilayering, where the cells are neither extruded from the tissue nor do they metastasize to distant organs even in presence of oncogenic signals^11–13^. Recently, the transcriptomic characterization of follicle cells with Lgl knockdown uncovered several co-existing heterogenous states of tumor-relevant gene expression that regulated the invasiveness of the ovarian multilayer^14^. Phenotypic characterization of these multilayered cells had previously revealed a distinguishable variability in their ploidy, that was found to get exacerbated when Notch signaling was ectopically induced (via N^ICD^ expression) in the Lgl-KD cells^12^. In this study, we have further investigated the underlying cause of this observable heterogeneity within the follicular multilayers.

The somatic epithelial cells of the *Drosophila* ovary emerge from dedicated stem cells, which are placed anteriorly within a structure called the germarium, to form polarized follicle cells that establish the monolayered epithelia which surrounds the germline cells within an egg chamber. During early oogenesis, while the polar and stalk-cell specification occurs at the earliest stages of follicle stem-cell differentiation, the remaining precursors form immature (and diploid) follicle cells that express Eyes absent (Eya) and divide via mitosis (in stage 1-6 egg chambers). These cells cease mitotic division and transition to recurrent endocycling at the onset of midoogenesis (stage 7) to form the mature (and polyploid) follicle cells expressing Hindsight (Hnt)^15^. Over the course of egg-chamber development, the monolayered epithelia are phenotypically molded by local microenvironmental cues, such as the Jak-STAT signaling ligand and mammalian IL-6 ortholog Unpaired (Upd), that are released from the polar cells (identified by the high expression of adhesion molecule Fas3) residing at the egg-chamber termini^13,16,17^. While the posterior follicle cells receive Upd from polar cells, that enables them to localize oocyte, anterior follicle cells adopt the border-, stretched- or centripetal-cell fate, respectively in response to Upd^18^. In this study, using single-cell RNA sequencing, we find that multilayered follicle cells activate a deviating transcriptomic response to the Upd signal, differing from their monolayered counterparts. Within the multilayer, Upd-exposed follicle cells exhibit phenotypes that are distinguishable from those beyond the reach of the Upd signal. In summary, we have identified an instance of ITH in a *Drosophila* tumor model that occurs in response to the microenvironment-derived Jak-STAT signal, by inducing one of the earliest characteristics of dysplastic cells, i.e., the loss of cell polarity.

## RESULTS

### Multilayered follicle cells exhibit heterogeneity of nuclear size and ploidy level

A normal egg chamber within the *Drosophila* ovary maintains a monolayered <u>f</u>ollicular <u>e</u>pithelia (FE) surrounding the developing germline cells (Figure 1A). To induce controlled multilayering in the FE, a pan follicle-cell driver *traffic-jam* (*tj*)-*GAL4* was used together with the temperature-sensitive (TS) *GAL80* repressor^19^. Using this *tj-GAL4, GAL80^TS^* (*tj^TS^*) driver, RNAi-mediated knockdown of *lgl* (Lgl-KD) was induced for 3 days, which resulted in the formation of epithelial multilayers at the egg-chamber termini during midoogenesis (Figure 1B). This Lgl-KD multilayer was significantly enhanced when the intracellular domain of Notch (N^ICD^) was ectopically expressed to provide oncogenic drive to the Lgl-KD cells (Figure 1C). Within both Lgl-KD and Lgl-KD+N^ICD^ expressing FE, a heterogeneity was observed in the intensities of RFP (transgenic reporter) expression among the multilayered cells. Within the Lgl-KD+N^ICD^ multilayers, the low-RFP cells were consistently found to be present in the vicinity of the polar cells, that were identified by high levels of Fas3^20^. Consistent with our previous observation^12^, the nuclei of these low-RFP cells appeared distinctly smaller than those of the high-RFP cells in the multilayer. To quantify the general relationship between RFP-reporter expression and the nuclear area, we measured them both in the control *tj^TS^* follicle cells (n=115). These cells were sampled from egg chambers present at progressive stages of development, since follicle cells normally increase their nuclear area following the onset of midoogenesis^21^. A positive correlation was indeed detected between the distribution of RFP intensities and the progressively-increasing developmental stages of follicle cells (Figure 1D and Supplementary Figure S1). When RFP intensities were measured and plotted against the per-quartile distribution of nuclear area for both Lgl-KD (n=57) and Lgl-KD+N^ICD^ (n=363) multilayered cells, a similar correlation was detected (Figure 1E-F). These results therefore suggest that the intensity of fluorescence reporter is related to the nuclear area of a cell.

**Figure 1.**
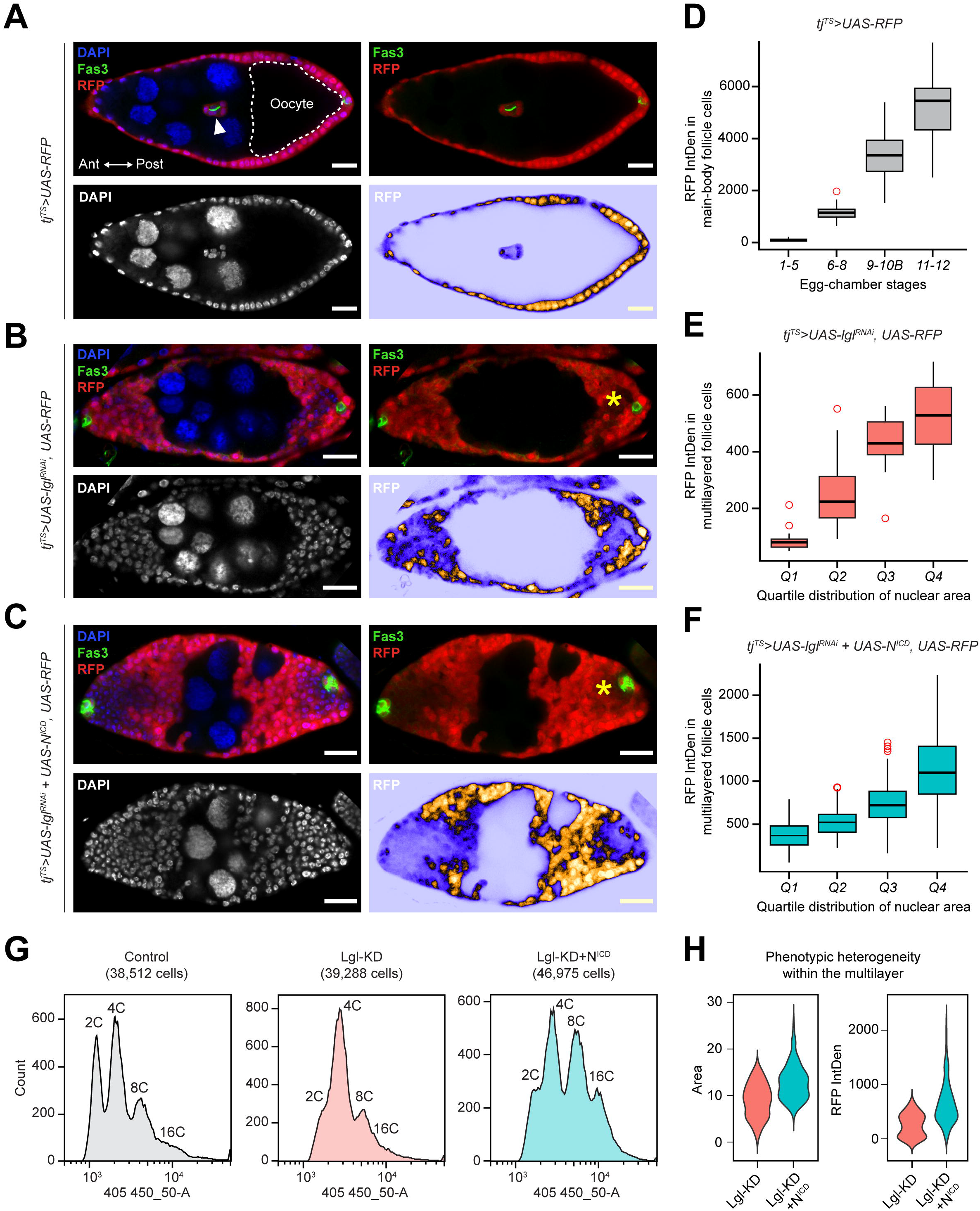
Multilayering of Lgl-KD expressing follicular epithelia exhibits intracellular heterogeneity. **(A)** Monolayered egg chambers containing RFP^+^ follicle cells (red) during midoogenesis follicle cells that form an epithelial monolayer around germline cells with larger nuclei (nurse cells) and the oocyte (dashed border). A pair of Fas3^+^ (green) polar cells appear within border cells (arrowhead) that delaminate from the anterior (Ant) epithelia, while another pair remains fixed in the posterior (Post) epithelia. DAPI is colored in blue. The top-right panel shows the relative position of Fas3^+^ polar cells and RFP^+^ follicle cells. Bottom panels specifically show DAPI (Left) and RFP intensities (Right). Comparable stages of egg chambers containing **(B)** Lgl-KD and **(C)** Lgl-KD+N^ICD^ in follicle cells are also shown. Follicle cells expressing uneven fluorescence intensities are indicated by asterisks (*). Scale bars: 20μm. **(D)** Box-and-whisker plot show the distribution of RFP intensity per cell (Y-axis) in *tj^TS^>RFP* control follicle cells at different stages of oogenesis (X-axis). Similar box-and-whisker plots show the distribution of RFP intensities in **(E)** Lgl-KD and **(F)** Lgl-KD+N^ICD^ cells within the multilayers (Y-axis) over increasing nuclear area that is binned into quartile distributions (X-axis). **(G)** Histograms show ploidy distribution of *tj^TS^-RFP* control, Lgl-KD and Lgl-KD+N^ICD^ follicle cells in respective samples. **(H)** Violin plots show the distributions of nuclear area (Left) and RFP intensities (Right) of each cell within the multilayered Lgl-KD and Lgl-KD+N^ICD^ follicle cells. See also Figure S1-S2.

Because the increase in nuclear area of follicle cells is a direct consequence of developmentally-regulated polyploidization over the course of egg-chamber development^22^, we wondered how the induction of Lgl-KD and Lgl-KD+N^ICD^ in follicle cells would affect the ploidy distribution of the developing FE. We used flow cytometry to isolate *tj^TS^*-control, Lgl-KD and Lgl-KD+N^ICD^ follicle cells and compared the distribution of cells with increasing ploidy. While a distinct peak separation was observed between the diploid (2C) and polyploid (>=4C) follicle cells in the experimental control, the “valley” separating the distinct 2C and 4C peaks was lost in the Lgl-KD sample, indicative of either a unimodal or a multimodal distribution of intermediate ploidies (Figure 1G). Similar loss of the 2C/4C peak separation was observed for Lgl-KD+N^IC^D follicle cells, where ectopic Notch activation is known to promote follicle-cell polyploidization independent of Lgl-KD^12^. A multimodal distribution was detected in the Lgl-KD+N^ICD^ egg chambers, as peaks were observed for both 2C and 4C ploidies alongside those attributable to likely intermediate ploidies populating the valley between them. Recently, polyploid cells in the *Drosophila* imaginal ring were shown to accelerate tumorigenesis by entering error-prone mitotic cell division and depolyploidization that result in increased DNA damage and ploidy heterogeneity^23^. The Lgl-KD+N^ICD^ follicle cells in the FE multilayers being polyploid as well, we sought to detect similar instances of DNA damage by using an antibody against γ-His2Av, that specifically identifies DNA double-strand breaks^24^. Enrichment of γ-His2Av foci was detected in the nuclei of the low-RFP expressing Lgl-KD+N^ICD^ cells within the multilayers, while the high-RFP cells did not display γ-His2Av foci (Supplementary Figure S2). This variable occurrence of DNA damage within the multilayers provides a likely explanation for the occurrence of intermediate ploidies that further promotes ITH within the growing neoplasm.

To summarize the observed ITH within both Lgl-KD and Lgl-KD+N^ICD^ multilayers, the distribution of nuclear area and RFP intensity in multilayered cells was assessed (Figure 1H). From the resulting violin plots, we concluded that: (1) given the multimodal ploidy distribution within the multilayer, the distinguishable phenotypes represent 2 or more populations within the multilayer, and (2) the Lgl-KD+N^ICD^ multilayer shows a greater range of variability. For these reasons, we then sought to determine the underlying differences between the Lgl-KD multilayer and that containing the Lgl-KD+N^ICD^ follicle cells with the intention of further identifying the underlying factors promoting the observable ITH.

### Multilayer heterogeneity associates with polar-cell derived paracrine Jak-STAT signaling

To further characterize the ITH in Lgl-KD+N^ICD^ multilayers, we performed whole-tissue RNA-sequencing (RNA-Seq) of ovaries containing Lgl-KD and Lgl-KD+N^ICD^ follicle cells alongside the follicle cells from the tj^TS^ experimental control that had been kept at 29°C to allow transgene expression for short (24h) or long (96h) periods of time (Supplementary Figure S3). Comparing the transcriptomes of all samples using Principal Component Analysis (PCA), we found that the only samples to form multilayers – Lgl-KD (96h) and Lgl-KD+N^ICD^ (96h) – clustered within the same quadrant separated from the rest (Figure 2A). This clustering reflected the overall transcriptomic similarity between the Lgl-KD and Lgl-KD+N^ICD^ multilayers, which is likely due to the mild autonomous upregulation of Notch (N) in Lgl-KD cells (0.76 log_2_ fold change (LFC) compared with the *tj^TS^* Control, *p*=0.03) (Supplementary Data 1). Further comparison of the Lgl-KD+N^ICD^ (96h) and Lgl-KD (96h) datasets revealed 412 genes differentially upregulated in the former, including those indicative of upregulated Jak-STAT signaling (*upd1* (4.3 LFC; *p*=1.9e-09), *upd3* (4.5 LFC; *p*=6.02e-08) and *Socs36E* (1.6 LFC; *p*=3.73e-06)). Markers of polar- and immature follicle-cell fates such as *Fas3* (1.26 LFC; *p*=3.35e-05), *castor* (*cas*) (1.96 LFC; *p*=0.02) and *eya* (1.3 LFC; *p*=4.06e-07) were also found upregulated in the Lgl-KD+N^ICD^ (96h) sample (Figure 2B and Supplementary Data 1). Consistent with these findings, we noticed a significant increase in the number of Fas3-expressing polar cells in the Lgl-KD+N^ICD^ multilayers (Figure 2C). From previous studies, N^ICD^ overexpression in follicle-cell precursors induces increased specification of polar cells^17^, which are the only cells within the FE that produce Jak-STAT ligand Upd1^25^. Therefore, the observed upregulation of Jak-STAT signaling in the Lgl-KD+N^ICD^ (96h) sample is a likely consequence of the increased number of polar cells.

**Figure 2.**
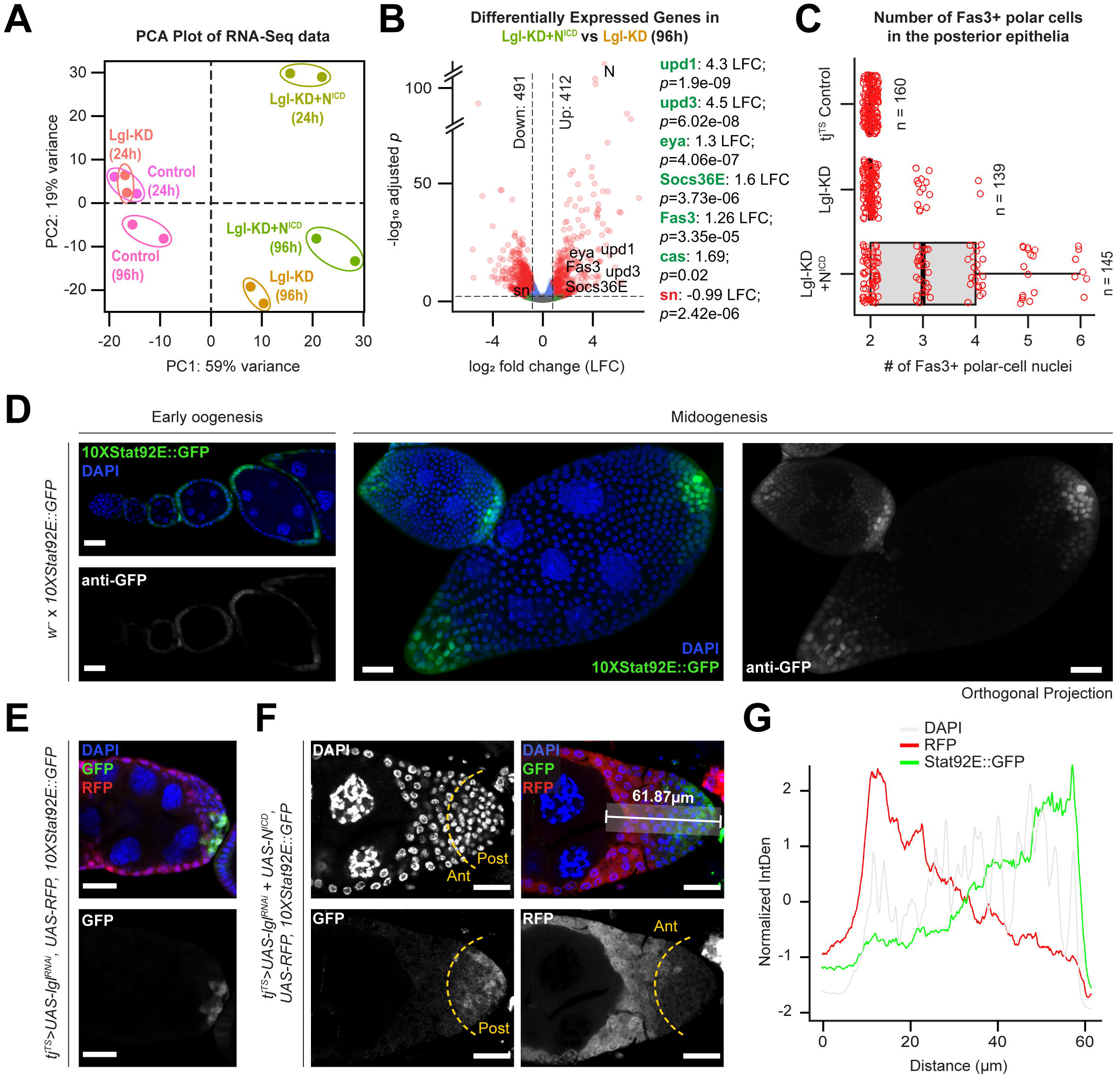
Multilayer heterogeneity is correlated with polar-cell derived Jak-STAT signaling. **(A)** PCA distribution of whole-tissue derived transcriptomes of ovaries containing follicle cells expressing *tj^TS^*-driven Lgl-KD and Lgl-KD+N^ICD^ for short (24h) and long (96h) time periods along with their corresponding control samples. Individual samples are uniquely colored and replicates per sample are encircled. **(B)** Volcano plot to show differential gene expression (dotted-line thresholds: > 0.4 log2 fold-change (LFC) and >= 0.05 p-value) in Lgl-KD+N^IC^D (96h) vs Lgl-KD (96h). LFC and p-values for upregulated (green) markers of polar cells and immature cells as well as genes associated with endogenous Jak-STAT signaling are shown along with that of the downregulated (red) border-cell marker expression. **(C)** Box-and-whisker plot to show the number of Fas3^+^ polar cells in the posterior egg-chamber epithelia containing follicle cells of different genotypes. 10XStat92E:GFP reporter expression (green) is shown in **(D)** *w*^-^ egg chambers at early and mid-oogenesis, as well as in the multilayers containing **(E)** Lgl-KD and **(F)** Lgl-KD+N^ICD^ follicle cells (red). The dotted yellow line arbitrarily separates the multilayer into the anterior (Ant) and posterior (Post) halves based on RFP expression. A 61.87μm long distance along the Ant-Post axis within the multilayer is highlighted in the top-right panel. The relative intensities (Y-axis) of normalized 10XStat92E:GFP (green), RFP (red) and DAPI (grey) expression over this distance is shown as a line graph **(G)**. Nuclei is marked by DAPI (blue). Scale bars: 20μm. See also Figure S3-S4.

Detection of FE-specific transcriptional profile from bulk RNA-Seq data is liable to contamination from ovarian cell types of non-epithelial origins. To circumvent this issue, we took advantage of the single-cell (sc) RNA-Seq datasets of ovaries containing *w^1118^* ^26^, Lgl-KD^14^ and Lgl-KD+N^ICD 12^ follicle cells to enable FE-focused analysis (Supplementary Figure S4). Using *w^1118^* follicle cells as reference, we integrated all 3 datasets and grouped the integrated cells into 11 clusters, that were roughly annotated using cell-type specific markers (Supplementary Figure S4). When the average expression of Jak-STAT pathway components was evaluated in each cluster, we found that the polar cells were the only ligand-producing cells in all 3 datasets, while the Jak-STAT signaling effectors were enhanced in the expected cells that receive them. Expression of the Jak-STAT pathway-driven transcription factors *ken, Stat92E* and pathway regulator *Su(var)2-10*^27^ were found to be elevated in (1) early-stage somatic cells in the germarium, (2) mitotic follicle cells, and (3) post-mitotic main-body follicle cells of the Lgl-KD+N^ICD^ dataset, when compared with cells of equivalent clusters in other datasets. These findings further support our conclusion from the bulk RNA-Seq analysis that Jak-STAT signaling was upregulated, and that its activation in Lgl-KD+N^ICD^ follicle cells was induced by the polar cells.

### Jak-STAT signaling maintains multilayered cells at developmental immaturity

Given the consistent occurrence of low-RFP cells adjacent to the polar cells, we argued that these cells were likely exposed to polar-cell derived Upd1 (or simply, Upd) in the microenvironment. Therefore, to verify Jak-STAT activation in the multilayer, we used the pathway reporter 10XStat92E-GFP^28^ that is normally detected in the terminal regions of the egg chambers containing monolayered FE (Figure 2D). Consistent with its normal expression as a gradient in posterior FE, the 10XStat92E-GFP reporter was detected in both Lgl-KD and Lgl-KD+N^ICD^ multilayers, with higher intensities of reporter expression in the posterior cells within the multilayer while little to no reporter activity was detected in the cells placed more anteriorly (Figure 2E-F). To further establish a link between pathway activation and ITH within the multilayers, we compared the normalized intensities of RFP and 10XStat92E-GFP in an area spanning 15 μm wide and 61.87 μm long over the posterior Lgl-KD+N^ICD^ multilayer shown in Figure 2F. An inversely proportional relationship between RFP intensity and 10XStat92E-GFP expression was observed within the multilayered FE (Figure 2G). As RFP was used as a proxy to distinguish ITH within the multilayer, this result confirmed that the cells displaying low RFP expression within the Lgl-KD+N^ICD^ multilayer exhibit differential Jak-STAT signaling activity.

As incremental changes in ploidy define the developmental transition from immature (mitotic) to mature (post-mitotic) follicle-cell lineage, we wondered if the multilayered cells with smaller nuclei exhibit characteristics of developmental immaturity. Using an antibody against the immature-cell marker Eya^29^, we found that Eya expression was restricted in follicle cells with low-RFP within the posterior multilayer of egg chambers during midoogenesis (Figure 3A). These Eya-expressing cells did not exhibit Cas expression (Supplementary Figure S1), a specific marker of the polar/stalk lineage^30^, which also partially overlaps in expression with Eya in the precursor follicle cells^30^, suggesting the Eya-positive cells in the multilayer are immature follicle cells. Next, we asked whether the cell-fate heterogeneity was induced by the restricted release of Upd in the microenvironment. To test this, we expanded the production of Upd to the entire FE by ectopically expressing Upd in Lgl-KD follicle cells, which induced unrestricted multilayering over the entire FE. All of these Lgl-KD+Upd-OE follicle cells showed Eya expression, while displaying low RFP intensity during midoogenesis (Figure 3A). These findings suggest that exposure of Lgl-KD cells to the Upd signal is key to their maintenance in an immature fate, and that the space restriction of the microenvironmental Upd signal promotes ITH in cell ploidy and cell fates.

**Figure 3.**
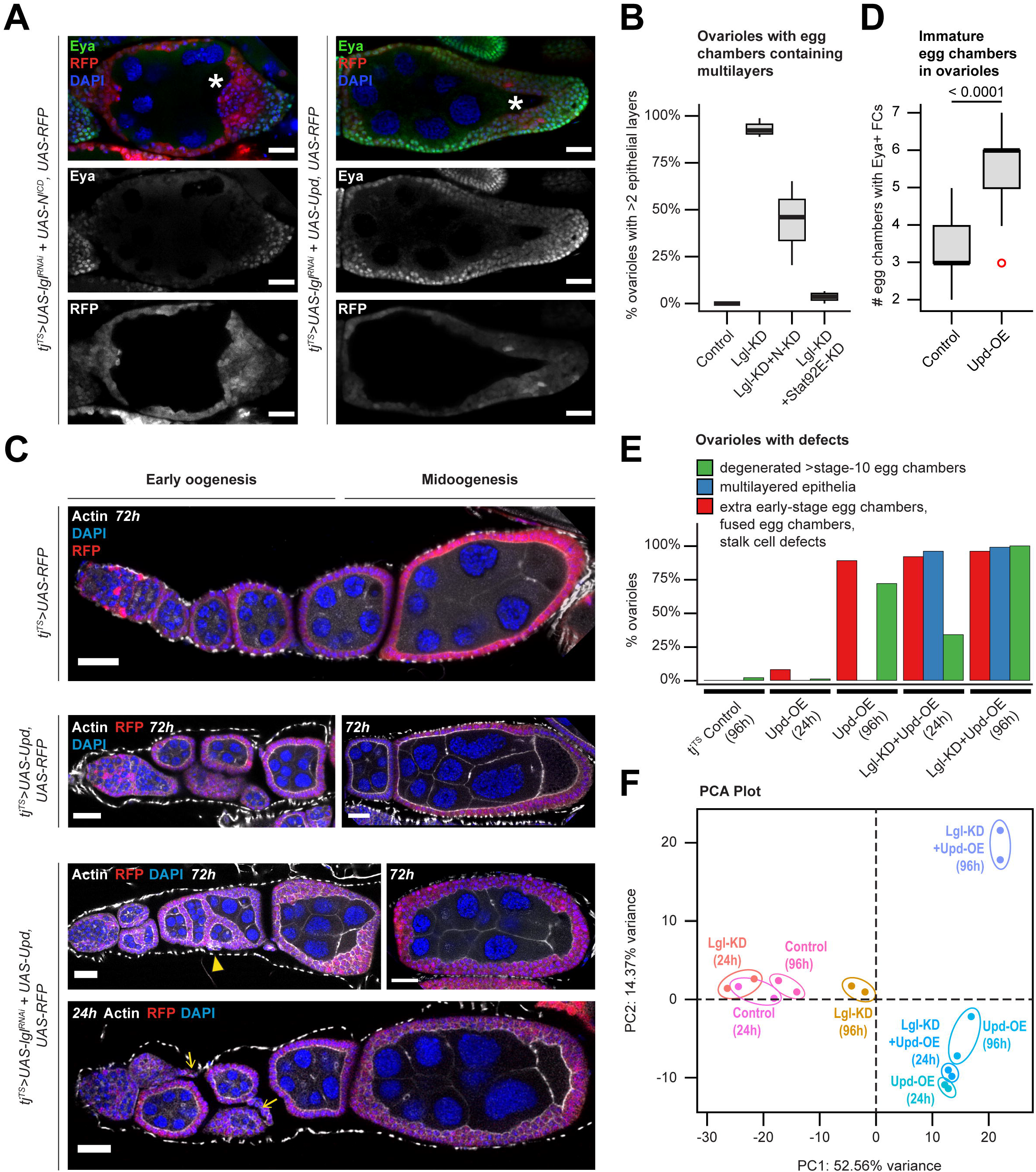
Unrestricted Upd-exposure induces defects in both normal and Lgl-KD follicle cells. **(A)** Confocal images of stage-8 egg chambers containing RFP^+^ (red) follicle cells with Lgl-KD+N^ICD^ (Left) and Lgl-KD+Upd-OE (Right) expression. Eya expression is detected within immature cells (green). Oocyte are marked by asterisks (*). Intensities of RFP expression and Eya antibody enrichment are shown separately (grayscale) in the middle and bottom panels. **(B)** Box-and-whisker plot to show the distribution of ovarioles from different genotypes (X-axis) with complete or partial (<2 layers) rescue of multilayering over 3 independent experiments. **(C)** Confocal images of egg chambers containing follicle cells expressing *tj^TS^*-driven RFP control (1^st^ row), Upd-OE for 72h (2^nd^ row), and Lgl-KD+Upd-OE for 72h (3^rd^ row) and 24h (4^th^ row) during early and midoogenesis. Abnormal stalk cells are indicated by yellow arrows while fused egg chambers with encapsulation errors are indicated by yellow arrowheads. Nuclei is marked by DAPI (blue). F-Actin is marked by Phalloidin (white). Scale bars: 20μm. **(D)** Box-and-whisker plot to show the number of egg chambers containing immature (Eya+) follicle cells (Y-axis) in individual ovariole, for when follicle cells express Upd-OE (compared with control). The p-value obtained from unpaired t-test comparison of the two distributions is indicated above the plot. **(E)** Bar plot to show the occurrence (%) of stage-specific ovarian phenotypes when Upd-OE and Lgl-KD+Upd-OE are expressed in follicle cells for 24h and 96h (compared with ovarioles with *tj^TS^* control follicle cells at 96h). **(F)** PCA distribution of whole-tissue transcriptomes of ovaries containing follicle cells expressing *tj^TS^*-driven Lgl-KD, Upd-OE and Lgl-KD+Upd-OE for short (24h) and long (96h) time-points, along with the corresponding control samples. Individual samples are uniquely colored and replicates per sample are encircled. See also Figure S5.

Because ectopic Upd or N^ICD^ expression regulates Lgl-KD multilayering by elevating Jak-STAT signaling either directly (via pathway activation) or indirectly (by increasing the number of signal-sending polar cells), we sought to determine how these manipulations differentially affect the heterogenous cell populations within the multilayer. We induced the knockdown of pathway effectors Notch (N-KD) and Stat92E (Stat92E-KD) individually in the Lgl-KD cells and assessed the resultant multilayered growth (Supplementary Figure S5). Comparison of the frequency of multilayer formation revealed a significant alleviation of the phenotype in Lgl-KD+Stat92E-KD egg chambers (3.87% ± 2.86 SD; n=120) from that of the Lgl-KD egg chambers (93.29% ± 4.98 Standard Deviation (SD); n=125), indicating an essential requirement of Jak-STAT signaling in promoting Lgl-KD multilayering (Figure 3B), as has been previously shown in the wing imaginal disc and the salivary gland imaginal ring^31^. In contrast, expressing N-KD in Lgl-KD cells showed a highly-variable rescue of the multilayer formation (44.16% ± 22.28 SD; n=124), as the number of cells expressing high RFP within the multilayer was significantly reduced, while the Eya-expressing, low-RFP cells continued to persist. These Eya-expressing cells did not get extruded from the tissue and continued to accumulate on the basal side of the FE without any detectable preference for dorsal or ventral positioning. Results from these experiments, therefore, further reinforced the conclusion that Jak-STAT signaling exclusively regulates and maintains the multilayered cells at developmental immaturity.

### Microenvironment-derived Upd elicits a novel response in cells with polarity loss

In the normally developing FE, posterior follicle cells respond to a gradient of paracrine Upd signal released from the polar cells with the purpose of specifying oocyte localization and egg-chamber polarization during midoogenesis^18^. Given that inducing polarity loss in the FE results in a dramatically changed phenotype, we wondered if cell-polarity loss would cause a cell to respond differently to similar microenvironment-derived cues. To distinguish between changes specific to Lgl-KD and Upd-OE in the FE, we sought to characterize the various ovarian phenotypes by separating them into stage-specific observations. Besides the expected developmental defects that manifest upon expressing Upd-OE alone in follicle cells, such as stalk-cell elongation and egg-chamber fusion^32–34^, we noted a statistically-significant increase in the number of egg chambers containing Eya-expressing cells within a single ovariole (5.51 ± 1.22 SD; n=227), from the control ovarioles during early oogenesis (3.11 ± 0.69; n=236) (Figure 3C-D). To analyze the phenotypes of egg chambers containing Lgl-KD+Upd-OE follicle cells, we used this “extra early-stage egg chambers” phenotype to distinguish the individual effects of Upd-OE and Lgl-KD. Using this guiding principle, the phenotypic changes associated with the expression of Lgl-KD and Upd-OE were quantified after allowing transgene expression for short (24h) and long (96h) periods of time (Figure 3D). The frequency of egg chambers containing Eya-expressing follicle cells within individual ovarioles increased upon sustained Upd-OE, with only 9.8% ovarioles (n=10/102) exhibiting the phenotype at 24h to 83.3% (n=105/126) at 96h. In contrast, ovarioles with Lgl-KD follicle cells showed a comparable extent of the extra egg-chamber phenotype after 24h (n=135/152) and 96h (n=108/118) of Upd-OE. Additionally, while Lgl-KD+Upd-OE expression for 96h caused unrestricted multilayering of the entire FE (n=117/118), this phenotype was also observed in the egg chambers at similar developmental stages that were allowed just 24h of transgene expression (n=147/152) (Figure 3E and bottom panel in Figure 3C). Quantifying the stage-specific phenotypes at early and late time points thus enabled the separation of the phenotypes that were induced by either Upd-OE or Lgl-KD. From our results, we concluded that the abnormal number of egg chambers during early oogenesis was exclusively caused by Upd-OE, while Lgl-KD was required for multilayer formation during midoogenesis.

We next sought to compare the whole-tissue transcriptomes of egg chambers that contained Upd-OE and Lgl-KD+Upd-OE follicle cells with those expressing Lgl-KD (as well as the experimental control) for short (24h) and long (96h) periods of time. When these datasets were disseminated on a PCA plot, Upd-OE was found to introduce the greatest variation (52.56% variance) in ovaries expressing just Upd in the FE. The two Upd-OE (24h and 96h) datasets were clustered together with the Lgl-KD+Upd-OE (24h) dataset in the same quadrant, indicating that at an early time-point, the gene expression state caused by Lgl-KD+Upd-OE is inseparable from that of Upd-OE alone. By contrast, the transcriptome of the Lgl-KD+Upd-OE (96h) sample was found separated from those of other Upd-OE samples along PC2 (14.37% variance), thereby suggesting that cells with polarity loss elicit a different response to microenvironment-derived Upd.

### Escargot is upregulated in follicle cells with Lgl knockdown and Upd overexpression

To further characterize how Lgl-KD follicle cells respond to Upd-driven Jak-STAT activation, we used scRNA-Seq to specifically identify the underlying gene expression of follicle cells. Generating the scRNA-Seq dataset for ovaries expressing Lgl-KD+Upd-OE in their follicle cells (for 72h) resulted in a total of 42,235 cells, where the germline cells, muscle sheath cells and hemocytes formed distinct clusters, but the follicle-cell lineage remained closely embedded (Figure 4A). Using specific markers (*bru1* and *osk* for early and late germline cells, *wupA* for muscle sheath, *Hml* for hemocytes, as well as *Sgs7* to identify a contaminant cluster of salivary-gland cells), the non-epithelial clusters were identified and subsequently removed from the dataset to obtain the remaining follicle cells. With the goal of characterizing the regulatory landscape within Lgl-KD+Upd-OE follicle cells, we identified their underlying regulon activity using SCENIC^35^. Among the binding motifs of TFs that control the identified regulons, the TPA-responsive element (TRE) sequence 5’-TGA/TCA-3’ and the E-box binding sequence 5’-CA(C/G)GTG-3’ were highly represented in the Transcription Start Sites (TSS) of genes expressed in the Lgl-KD+Upd-OE cells (Figure 4C). TRE-binding families of TF such as AP-1, CREB and ATF^36^ were recently detected in follicle cells with polarity loss via Lgl-KD^14^. The Ebox binding motif is recognized by the bHLH-Zip family of TFs such as Myc, Max, Emc etc.^37^ as well as those belonging to the SNAIL family TFs, such as Snail (Sna), Escargot (Esg) and Worniu (Wor)^38^. None of these TF-associated regulons showed cluster-specific activity in the scRNA-Seq dataset, thus indicating that they were active globally in all Lgl-KD+Upd-OE cells.

**Figure 4.**
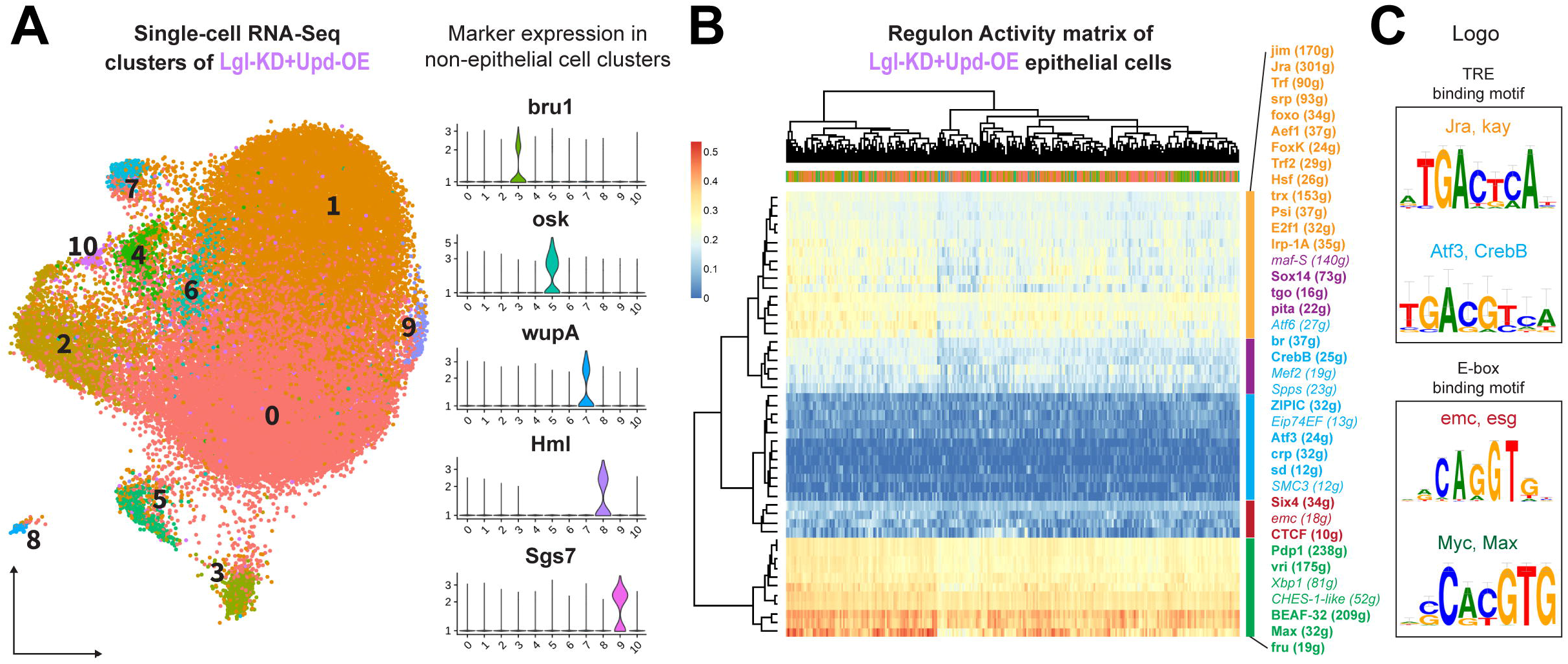
scRNA-Seq identifies regulon activity within Lgl-KD+Upd-OE follicle cells. **(A)** UMAP plot to show the 42,235 ovarian-cell types grouped into 11 clusters that are identified by their unique colors and ID numbers embedded on the plot. Adjacent violin plots identify the relative expression (Y-axis) of non-epithelial cell markers that are expressed in specific clusters (X-axis). **(B)** Heatmap to show the activity of regulons (rows) in a random subset of 300 cells (columns) from each epithelial-cell cluster (i.e., clusters 0-2, 4, 6, 10). Both rows and columns are hierarchically clustered and the tree is shown. Individual regulons are identified by the regulating transcription factor (TF) while genes within each regulon is mentioned within parentheses next to the TFs. The regulons are colored according to their cluster IDs, which are split into the top 5 clusters. Italicized regulons include low-confidence associations. **(C)** UMAP plot of 42,235 single cells from Lgl-KD+Upd-OE (72h) dataset, grouped into 11 clusters. Violin plots (Right) showing cluster-specific expression of non-epithelial cell markers. **(D)** DNA-binding motifs related to TRE and E-box sequences are shown that correspond to the active regulons that are indicated above each logo. See also Figure S6.

We attempted to identify the cell types in the Lgl-KD+Upd-OE dataset by integrating it along with the annotated datasets of *w^1118^*, Lgl-KD and Lgl-KD+N^ICD^ follicle cells^12,14,26^. A total of 76,763 follicle cells from all datasets were integrated and grouped into 21 clusters that were then roughly annotated for cell-type identities using the expression of known markers (Figure 5A; Supplementary Figure S6). This dataset integration revealed a significant deviation of the Lgl-KD+Upd-OE follicle cells from the expected follicular lineage, as most of them were found to populate clusters 0, 1, 2 and 5 that overlapped with cells from the Lgl-KD and Lgl-KD+N^ICD^ datasets (Figure 5B-C). Expression of Esg (the *Drosophila* ortholog of Snail2) was detected specifically in Lgl-KD+Upd-OE cells of those clusters, unlike that of Emc and Eya that are detected in all immature (Cut expressing) cells and in some mature (Hnt expressing) cells of the normal follicle-cell lineage^39^ (Figure 5D). We sought to validate the expression of Esg in the heterogenous cells of the Lgl-KD+N^ICD^ multilayer by using a protein-trap construct that incorporates GFP in-frame at Esg C-terminus^40^. Using anti-GFP antibody to enhance detection of the lowly-expressed protein, Esg enrichment was found in the nuclei of the low-RFP cells within the posterior multilayer (Figure 5E). In summary, scRNA-Seq analysis identified the differential expression of Esg within Lgl-KD+Upd-OE follicle cells, and its expression was also validated in multilayered cells exposed to endogenous Upd.

**Figure 5.**
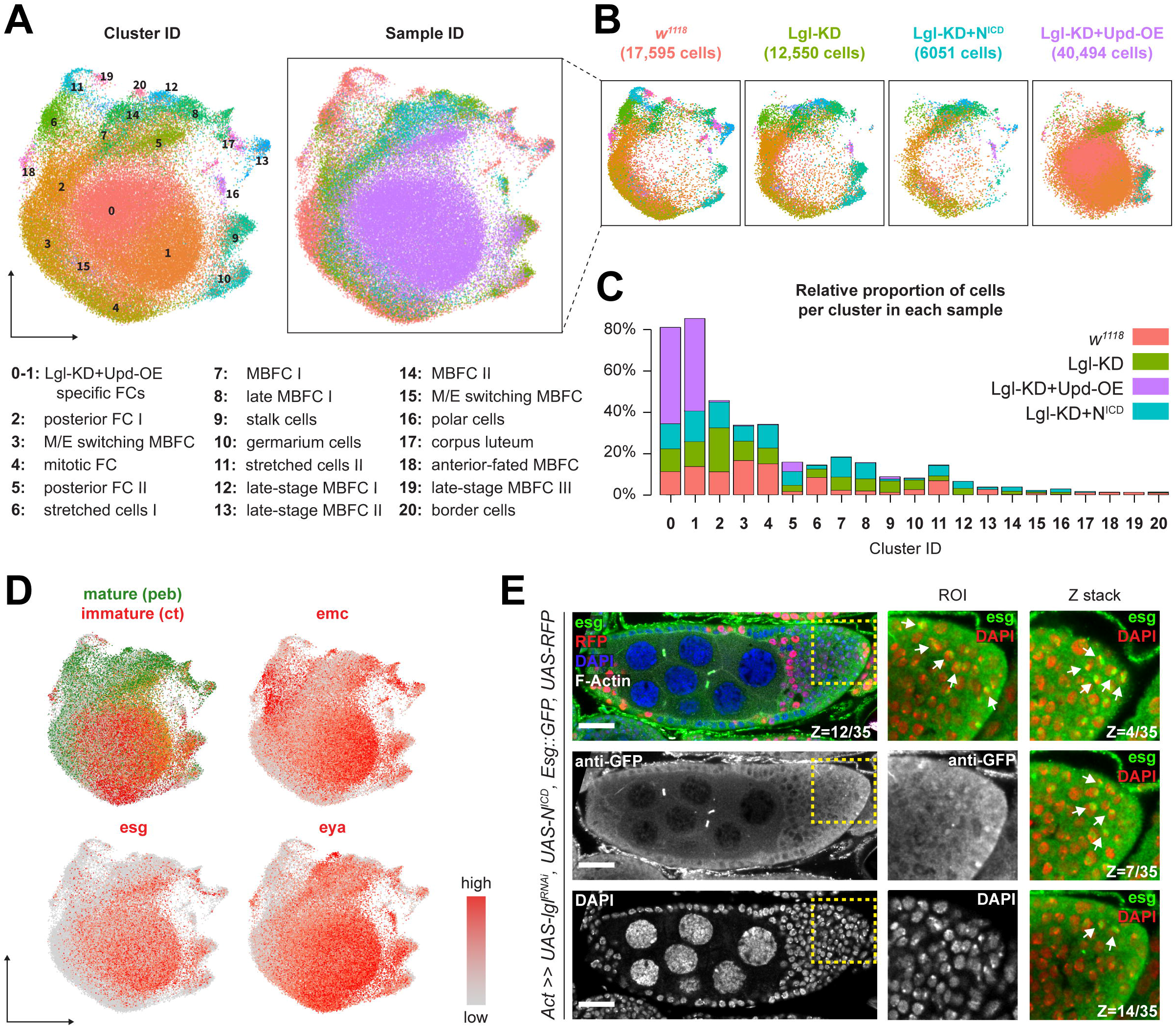
Esg activation is detected within Lgl-KD follicle cells exposed to Upd. **(A)** UMAP plot to show the 76,763 follicle cells of the integrated dataset, colored by their cluster ID (Left) and sample ID (Right). Estimated cell-type identities of individual clusters are indicated below. FC = Follicle Cells; MBFC = Main Body FCs. **(B)** UMAP plots split by sample ID, where individual cells are colored by their cluster ID. **(C)** Bar plot to show the proportion of cells colored by their sample ID (Y-axis) in each cluster (X-axis). **(D)** Enrichment plots to show the expression (red) of *emc*, *esg* and *eya* in the integrated follicle cells. Mature and immature follicle cells are identified by the differential enrichment of *peb* (green) and *ct* (red) respectively. **(E)** Confocal image of multilayered egg chambers containing RFP^+^ Lgl-KD+N^ICD^ follicle cells (red), where the protein localization of Esg is enhanced by anti-GFP antibody (green) within a 35μm thick (Z-axis) tissue (Left panel). Nuclear enrichment of Esg (marked by arrows) is shown separately for a Region of Interest (ROI) that is identified by a dotted, yellow square, with distinct panels to show the overlap of Esg (green) and DAPI (red), as well as Esg and DAPI staining alone (greyscale; middle column). Nuclear enrichment of Esg is also shown for cells at different sections along the Z-axis (right column). Scale bars: 20μm. See also Figure S6.

### Escargot drives cell-fate defects in Lgl-KD follicle cells in response to Jak-STAT signaling

Next, to test the possible role of Esg in Lgl-KD multilayers, we expressed Esg (Esg-OE) in Lgl-KD follicle cells and compared its phenotype with egg chambers containing follicle cells expressing Esg alone. While expressing Esg in follicle cells alone caused a 38.32-fold increase in Esg transcripts (assessed by quantitative Real-Time PCR or qRT-PCR), no discernable changes to egg-chamber development (compared with normal tissue) were detected (Figure 6A). In contrast, in egg chambers containing Lgl-KD+Esg-OE follicle cells, FE organization was found severely disrupted (Figure 6B). These egg chambers showed consistent mis-localization of the oocyte, egg-chamber fusion as well as a global differentiation failure of the follicle-cell lineage, as indicated by the widespread expression of Fas3, a polar-cell marker that is also expressed weakly in the precursor follicle cells within the germarium. Lgl-KD+Esg-OE follicle cells also displayed sustained expression of Eya (n=49/56 intact egg chambers) and absence of Hindsight (Hnt) (n=48/48), similar to follicle cells with Lgl-KD+Upd-OE (Figure 6C). Moreover, in a manner similar to that observed in Lgl-KD+Upd-OE follicle cells, both Eya and Hnt were found to correlate with RFP expression and nuclear area of these Lgl-KD+Esg-OE follicle cells, where high Eya was detected in low-RFP expressing cells with smaller nuclei and Hnt was only detected in high-RFP cells with larger nuclei (Figure 6D). These observations suggest that Esg expression phenocopies Upd in affecting the cell fate of Lgl-KD follicle cells.

**Figure 6.**
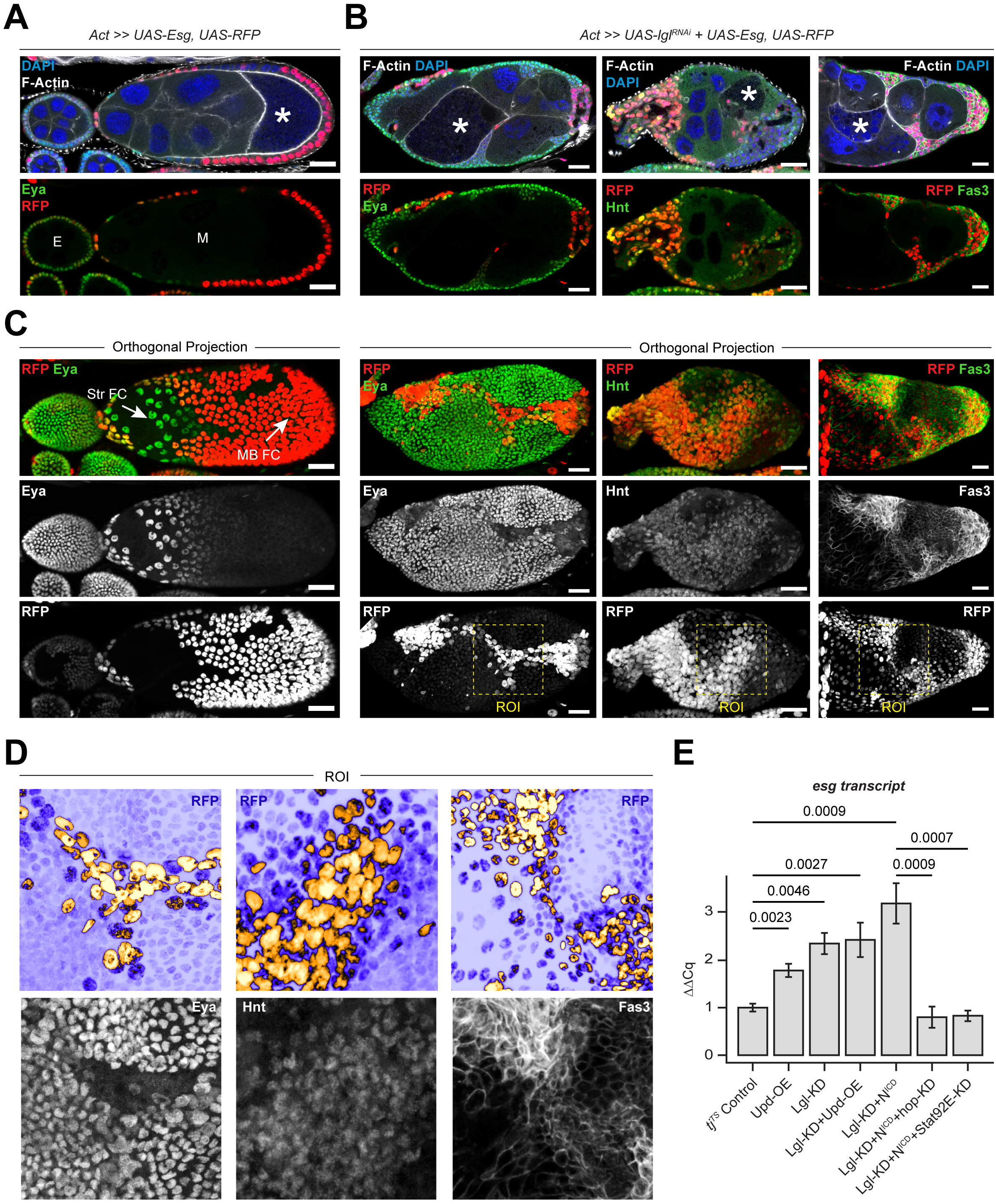
Esg induces cell-fate changes downstream of Lgl-KD and Jak-STAT signaling. **(A)** Confocal images of egg chambers with RFP+ (red) follicle cells ectopically expressing Esg. Eya staining (green) is used to mark undifferentiated during early (E) and stretched follicle cells (Str FC) during mid- (M) oogenesis. Main-body follicle cells (MB FC) express RFP at high intensities and lack Eya expression. Oocyte is indicated by asterisks (*). **(B)** Egg chambers at midoogenesis containing RFP^+^ Lgl-KD+Esg-OE follicle cells. Cell-fate defects are shown by the aberrant expression of Eya, Hnt and Fas3. Mis-localized oocytes are identified by asterisks (*). **(C)** Orthogonal projections of egg chambers shown in panels A-B. Overlap between RFP (red) and Eya, Hnt and Fas3 respectively (green) is shown. Each channel is also shown separately. A Region of Interest (ROI) is highlighted as the region within a yellow dotted square. Scale bars: 20μm. **(D)** Zoomed-in images of the ROIs in panel C. The top panels show RFP-expression intensity (yellow indicates high and purple indicates low intensity), while the bottom panels show Eya (Left), Hnt (middle) and Fas3 (Right) enrichment (greyscale). **(E)** Bar plot to show the qRT-PCR results, where Y-axis shows the relative normalized expression (ΔΔCq) of *esg* mRNA in the different genotypes (X-axis). P-values derived from t-tests for statistical significance between individual genotypes are mentioned on top.

We then examined if Esg functions downstream of Jak-STAT activation in the Lgl-KD cells. Using qRT-PCR, we assessed the levels of *esg* transcripts in whole-tissue samples containing follicle cells expressing transgenes under the *tj^TS^* driver. Compared with the experimental control, samples of ovaries containing Lgl-KD, Lgl-KD+Upd-OE, and Lgl-KD+N^ICD^ follicle cells showed 2.33 (± 0.23 Standard Error), 2.41 (± 0.2 SE) and 3.17-fold (± 0.24 SE) increase in Esg-transcript abundance, respectively. Inducing the knockdown of Hopscotch, the Jak ortholog in flies (Hop-KD) and Stat92E, the STAT ortholog (Stat92E-KD) in the Lgl-KD+N^ICD^ follicle cells then caused 0.79 (± 1.27 SE) and 0.83-fold (± 0.06 SE) reduction, respectively, in Esg expression compared to the control (Figure 6C). Given these results, we concluded that Esg is likely activated downstream of the Jak-STAT signaling cascade within Lgl-KD+N^ICD^ cells. To test this relationship further, we knocked down Esg expression (Esg-KD) in Lgl-KD follicle cells, where a decrease in the number of Eya-expressing cells was found within the multilayers, which were also significantly reduced in volume (Supplementary Figure S7). We then induced Esg-KD in Lgl-KD+Upd-OE follicle cells and evaluated for a rescue of the severe phenotype. To drive Esg-KD in Lgl-KD+Upd-OE background, we used the *Act5C*-promoter regulated *GAL4* driver to genetically accommodate all relevant transgenes. Inducing Lgl-KD+Upd-OE in FE thus induced severe phenotypic defects that caused egg chambers (n=16/23) to form completely disrupted structures with dying germline cells (Figures 6D). Few egg chambers (n=7/23) showed defects in germline-cell encapsulation that was likely caused by defects in follicle-cell differentiation, as shown by abnormal Eya and Hnt expression. Inducing Esg-KD in these cells significantly alleviated the severe phenotype, where the Upd-OE specific “extra egg-chamber” defect was observed during early stages (n=42/48) and a rescue of multilayering was observed during midoogenesis (n=15/42), where cells continued to express Eya. While ~64% of ovarioles (n=27/42) continued to exhibit both multilayered FE and germline-cell encapsulation, the phenotype was found to be partially reduced from that observed for the egg chambers with Lgl-KD+Upd-OE follicle cells. Taken together with our previous result where Esg expression is reduced upon the knockdown of the Jak-STAT signaling components, we concluded that Esg indeed functions downstream of Jak-STAT signaling activation in the Lgl-KD follicle cells. However, this conclusion is somewhat tempered by the incomplete rescue (by Esg-KD) of the severe Lgl-KD+Upd-OE phenotype, which is probably indicative of redundant processes that regulate one or more phenotypes that result from Lgl-KD+Upd-OE expression in the FE.

### Escargot is required during early follicle-cell differentiation

Intriguingly, overexpression of Upd in follicle cells alone resulted in a 1.73-fold (± 0.09 SE; *p*=0.0023) increase of *esg* mRNA, which is normally expressed in the Escort Cells (ECs) that lie anterior to FSCs and therefore being antecedent to the follicle-cell differentiation^41,42^. The increase in *esg* mRNA level prompted us to determine the origin of these transcripts in the ovary. We first queried the publicly-available datasets from two independent *Drosophila* ovarian-cell atlas resources^26,42^ and found that Esg was highly expressed in the germline-cell clusters and at significantly low levels in the somatic cells of the germarium (data not shown). Next, we used another publicly-available dataset that was generated with specific focus on the earliest stages of oogenesis^43^. Similar to our assessment from the aforementioned cell atlases, *esg* was detected at traceable amounts within the various EC clusters and at high levels in the clusters representing the early germline-cyst cells (Figure 7A). We validated the enrichment of Esg using the previously-mentioned protein-trap line, which detected the protein within the germline cysts immediately posterior to germline stem cells (GSCs) as well as in the oocytes of early-stage egg chambers (Figure 7B). In the somatic cells, in addition to its expression in the cone-shaped ECs that occupied region 2a of the germarium, Esg was detected in the FSCs at region 2b as well as in the follicle-cell lineage posterior to the FSCs. While it was technically not feasible to incorporate the protein-trap line with *tj^TS^*-driven Upd-OE, we also visualized Esg enrichment in early-stage follicle cells expressing N^ICD^ (which mimic Upd-OE by increased polar-cell specification as shown earlier), that presented a similar outcome as observed in the normally developing ovariole (Supplementary Figure S7). The upregulated *esg* transcripts in the ovaries with Upd-OE follicle cells could therefore be derived from both somatic and germline cells, as the germline cells also expands when the adjacent somatic cells overexpress Upd in the germarium^44,45^.

**Figure 7.**
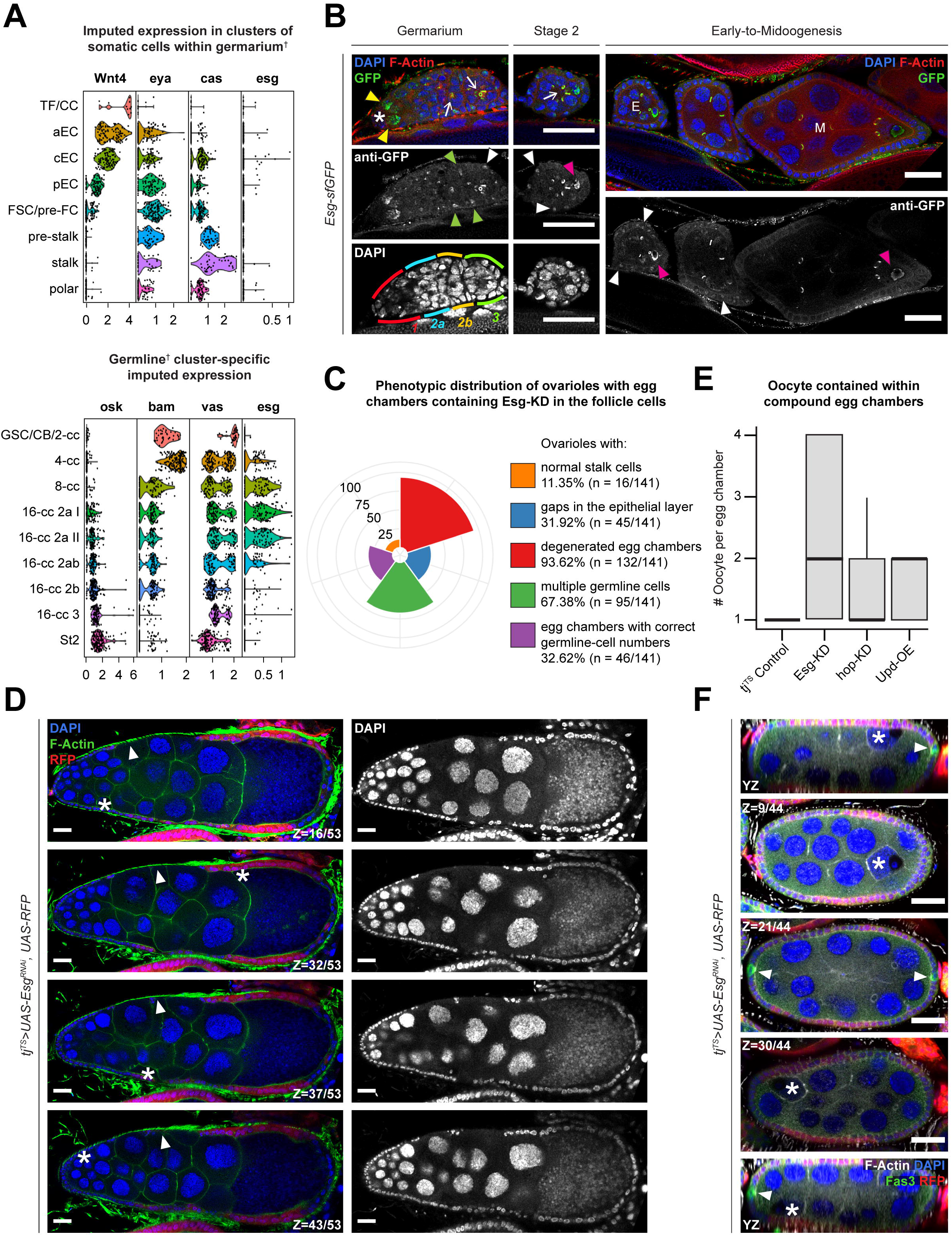
Escargot is required for proper egg-chamber development. **(A)** Confocal images showing the GFP-antibody enhanced nuclear detection of Esg protein (green) localization in early oogenesis. GSCs are indicated by asterisks (*), while Esg enrichment in early cyst cells are indicated by yellow arrowheads and oocytes are indicated by pink arrowheads. Ring canals connecting the differentiating germline cells are indicated by arrows. Esg enrichment in the FSCs are marked by green arrowheads and white arrowheads mark the immature follicle cells in early (E) egg chambers. Note, that Esg is not detected in follicle cells in egg chambers at midoogenesis (M). The various regions of the germarium is shown by distinct colors (red = region 1; blue = region 2a; yellow = region 2b; and green = region 3) as mentioned within the figure. Violin plot showing the imputed expression of *osk, bam, vasa* and *esg* in **(B)** germlinecell clusters and that of *Wnt4, eya, cas* and *esg* in **(C)** somatic cells within germarium in the Slaidina *et al*. scRNA-Seq dataset. ^†^Clusters are presented as determined by the original authors of the dataset. **(D)** Polar bar plot showing the relative percentage of specific (individually color-coded) phenotypes, as shown in figure legend to the right of the plot. **(E)** Box-and-whisker plot showing the number of oocytes (Y-axis) presented within intact egg chambers of different genotypes (X-axis). **(F)** Confocal image showing compound egg chambers formed by expressing Esg-KD in RFP^+^ (red) follicle cells. Individual oocytes shown at different sections on the Z-axis are marked by asterisks (*), while gaps in the FE are indicated by white arrowheads. **(G)** Confocal image showing a compound egg chamber highlighting Fas3^+^ (green) polar cells (arrowheads) and mis-localized oocytes are indicated by asterisks (*). Nuclei are stained with DAPI. F-Actin is stained with Phalloidin. Scale bars: 20μm. See also Figure S7.

To assess whether Esg expression in the somatic cells plays a role in follicle development, we used the *tj^TS^* driver to knockdown Esg in the FE, as well as the ECs and FSCs. In the resultant ovaries, only 11.35% (n=16/141) ovarioles showed normal morphology with stalk cells connecting individualized egg chambers during early oogenesis, while the rest showed no separation of egg chambers and an absence of stalk cells (Figure 7C-D). Gaps in the FE were observed in 31.92% of all ovarioles (n=45/141), whereas 93.62% (n=132/141) had dying germline cells at around stage 10. Compound egg chambers, a phenotype associated frequently with a failure in follicular encapsulation upon the chronic misexpression of Jak-STAT pathway components Upd and Hop^32,33^, were observed in 67.38% of all ovarioles (n=95/141) containing Esg-KD follicle cells. Inside these egg chambers containing Esg-KD FE, significantly higher numbers of oocytes were detected (2.22 ± 1.29 SD), compared to those containing Upd-OE (1.53 ± 0.5 SD) and hop-KD (1.53 ± 0.58 SD) respectively (Figure 7E). Additionally, Esg-knockdown induced compound egg chambers showed mislocalization of the oocytes—although they remained attached to the FE, they were no longer adjacent to the polar cells (Figure 7F). As oocyte localization depends on proper Lgl function in posterior follicle cells^11^ and their response to polar-cell derived Upd^18^, our results suggest that Esg expression in follicle cells is likely downstream of Jak-STAT signaling and necessary for follicle-cell differentiation.

## DISCUSSION

In this study, we introduced a novel model of induced heterogeneity, where loss of polarity in the *Drosophila* FE promotes heterogenous multilayering of cells that continue to grow *in situ*. As the multilayer volume expands, cells that remain exposed to Upd (fly ortholog of IL-6) in the microenvironment are maintained at developmental immaturity, thereby introducing cell-fate heterogeneity within the nascent neoplasm. Unlike the monolayered follicle cells at comparative stages of egg-chamber development, polarity-deficient immature cells activate the expression of Esg, (fly ortholog of Snail2, also known as Slug). Activation of Esg keeps the polarity-deficient Lgl-KD follicle cells at an immature fate, but does not affect the polarized follicle cells. Our findings indicate that Esg functions downstream of Jak-STAT signaling exclusively in the follicle cells that lack epithelial-cell polarity, a relationship that has been previously reported in the hyperplastic-eye tissue^46^. In humans, Cancer-Associated Fibroblasts (CAFs) are among the most abundant non-tumor cells within the tumor microenvironment^47^ that communicate with tumor cells via the IL-6/STAT3 signaling axis^48,49^, which has been found to regulate tumor-cell dedifferentiation and cancer stem cell maintenance^50^ via Snail^51,52^. Extrapolating our results to mammalian tumor cells that exhibit dissimilar loss of epithelial polarity depending on their location within the tumor tissue (cells at leading edge vs lagging cells) and their mode of invasion (amoeboid vs mesenchymal vs collective migration)^53,54^, exposure to permissive microenvironmental signals could cause diverse effects in each cell promoting phenotypic heterogeneity.

Independent of our findings in the FE tumor model, we also identified a comparable link between Esg and cell-polarity loss in the somatic cells of the germarium, where the Esg-expressing cells exhibit a remarkable lack of polarized architecture. These cells represent the Escort Cells (ECs) in region 1-2a, and the Follicle Stem Cells (FSCs) in region 2b of the germarium. Spanning regions 1 and 2a, the ECs are divided into the anterior, central and posterior domains, among which, the posterior ECs (pECs) exhibit lineage plasticity which allow them to exchange cell-fate identities with the FSCs^42,55,56^. This recently-uncovered interchangeability among the somatic-cell populations of the germarium, therefore, offers a likely explanation of why the translated product of Esg – which is otherwise expected to stay restricted to the ECs^41,42^ – is detected in both the FSCs and in the immature follicle cells. Given our finding that Esg expression is essential for normal FE development, while also recognizing that none of the epithelial cells at comparable developmental stages show enrichment of the *esg* mRNA, we believe that Esg knockdown adversely affected the pECs that convert to FSCs, thereby causing a domino effect in the subsequent progenies that make up the FE. We propose that within the germarium, Esg is activated in the polarity-deficient ECs when they’re exposed to Upd in region 2b, but its expression is immediately blocked upon the polarization of the follicle cells as they convert to FSCs, which would explain its low level of detection via RNA-Seq.

Jak-STAT signaling in the germarium has been linked with proliferation and fate maintenance of the pECs^56^ as well as proliferation of the immediate progenies of FSCs^44^. In our experiments, when Upd alone is overexpressed in the FE, extra egg chambers containing immature follicle cells were observed during early oogenesis, that infrequently showed encapsulation failures. In contrast to the studies where *C857-GAL4* was used to drive Upd in the somatic cells of the germarium that caused germline hyperproliferation^44,45^, we used the pan follicle-cell driver *tj-GAL4* to sustain Jak-STAT signaling activation in the FE beyond the confinement of the germarium. Given this setup, we argue that the increase in the numbers of egg chambers is likely caused by the combined increase in the proliferation rate of both germline and somatic follicle cells, which does not cause FE multilayering but instead exhibits the occasional failure of egg-chamber encapsulation. Thus, we have also uncovered a previously overlooked role of Jak-STAT signaling in egg-chamber development during early oogenesis.

While “unlocking phenotypic plasticity” is now acknowledged as an emerging hallmark of cancer cells^57^, the process is observed more commonly during normal development^58,59^. In this study, we have revealed significant differences between the plasticity induced by Upd in the normal FE and that seen in the multilayered cells where exposure to Upd induces cell-fate changes, ploidy heterogeneity and DNA double-strand breaks. Endocycling follicle cells have been shown to undergo depolyploidization and exhibit genomic instability when forced to reenter mitosis in a manner that is independent of fate regulation in these cells^60^. In our study, using RFP expression as a proxy for ploidy, we discovered that the Esg-expressing Lgl-KD follicle cells with intermediate ploidy levels were found in the transient zone separating the cells with higher and lower ploidies (Figure 6D). This observation therefore indicates that the polyploid follicle cells expressing Lgl-KD+Esg-OE likely undergo depolyploidization to produce cells with serially decreasing ploidies (and thus, RFP) to populate the FE. Instances of depolyploidization and error-prone mitosis are often considered to be the critical drivers of heterogeneity in a growing neoplasm^23,61–63^ and it could likely be playing a similar role within the FE multilayers. Thus, our results offer quite a few plausible explanations for how heterogeneity could be introduced within genetically-identical cells in response to selective pressures from the microenvironment.

### Limitations of the study

While Upd-mediated Esg activation in the polarity-deficient cells could sufficiently drive heterogeneity by forcing cells to remain immature, the alternative that heterogeneity could be caused by the asynchronous accumulation of cell-cycle arrested cells in the multilayer is equally likely. The resultant ploidy heterogeneity is expected to emphasize copy-number diversity within the multilayer that could further reinforce the observed heterogeneity and drive changes independent of Upd. Furthermore, even though a previous study showed that genomic instability in endocycling follicle cells that are forced to re-enter mitosis is independent of cell fate determination^60^, we were still able to use mature/immature follicle-cell markers to characterize the cell-fate heterogeneity in the multilayer. These defects in ploidy, therefore, likely represent the tissue-specific changes that are caused by using N^ICD^ as the oncogenic driver, which also drives cell-fate and cell-cycle switching in follicle cells. It is thus possible that some aspects of the described changes to the cells represent developmental defects that may not be recapitulated within the tumor as a general outcome of intra-tumor heterogeneity. While these tissue-intrinsic properties somewhat temper our conclusions, the central idea presented in this study offer a testable hypothesis linking polarity loss to Snail2 activation in mammalian neoplasms in response to Jak-STAT signaling, both being common properties of cancer cells.

## ACKNOWLEDGEMENTS

We thank Allison Jevitt, Gengqiang Xie, Su-Mei Zhang, Cynthia Vied, Amber Brown, and Brian Washburn (at Florida State University Biological-Science Core) for their assistance in library preparation and sequencing. 10X Chromium controller and other essential hardware was provided by the Florida State University Translational Science Laboratory, while the server space for computational analyses were provided on the Cypress server system by Tulane University. We would also like to thank Chun-Ming Lai, Megan Rasmussen, Virginia Morejon and other Deng lab members for helpful discussion and manuscript preparation. Finally, Bio-Render (licensed by Tulane University) was used to generate images for the Graphical Abstract.

## AUTHOR CONTRIBUTIONS

Conceptualization, D.C. and W.D.; Methodology, D.C.; Software, D.C.; Validation, D.C. and C.A.M.C.; Formal Analysis, D.C.; Investigation, D.C., F.C., C.A.M.C. and X.F.W; Resources, Y.C.H.; Data Curation, D.C.; Writing – Original Draft, D.C.; Writing – Review & Editing, D.C., W.D.; Visualization, D.C.; Supervision, W.D.; Project Administration, W.D.; Funding Acquisition, W.D.

## STAR METHODS

### KEY RESOURCES TABLE

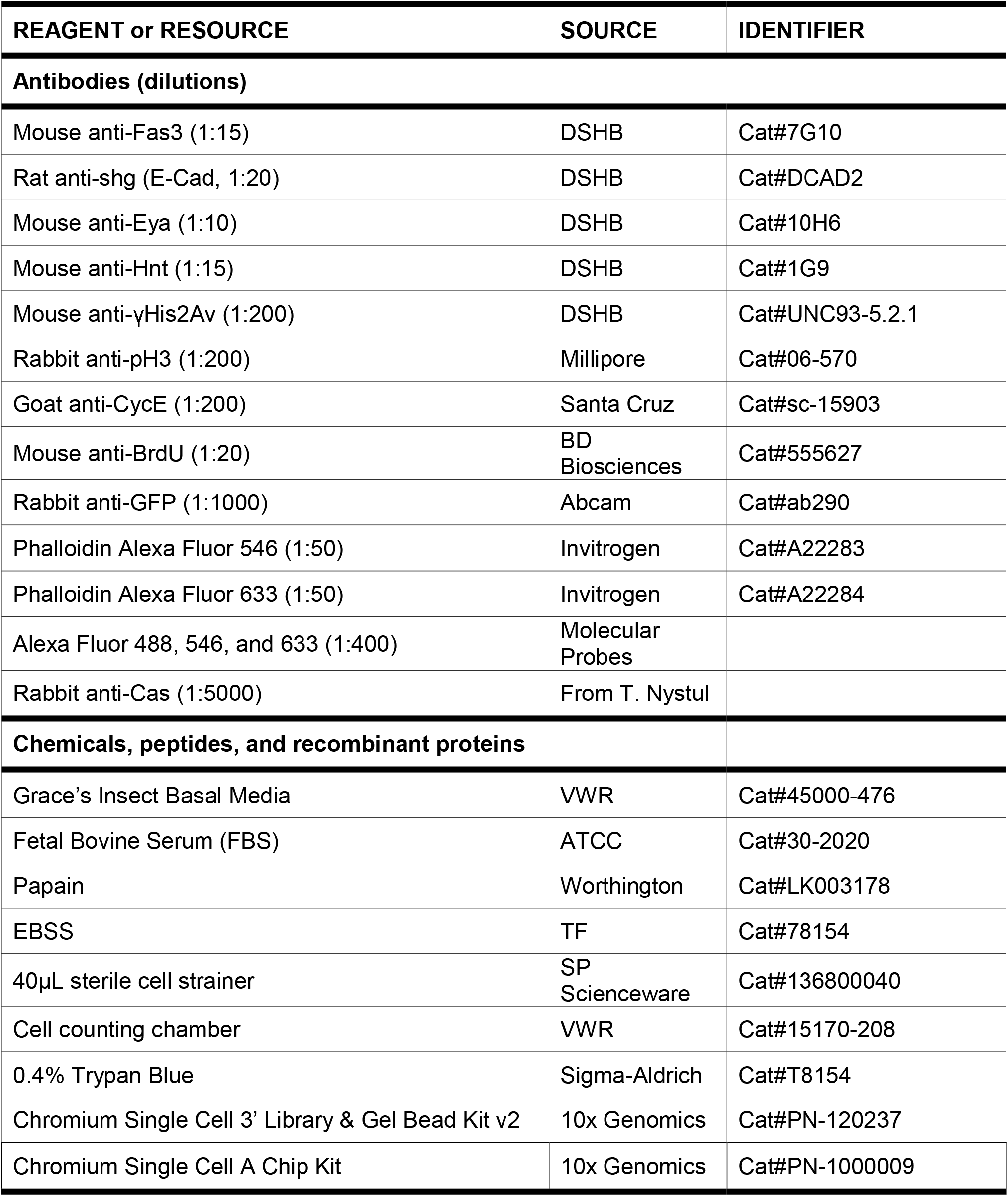
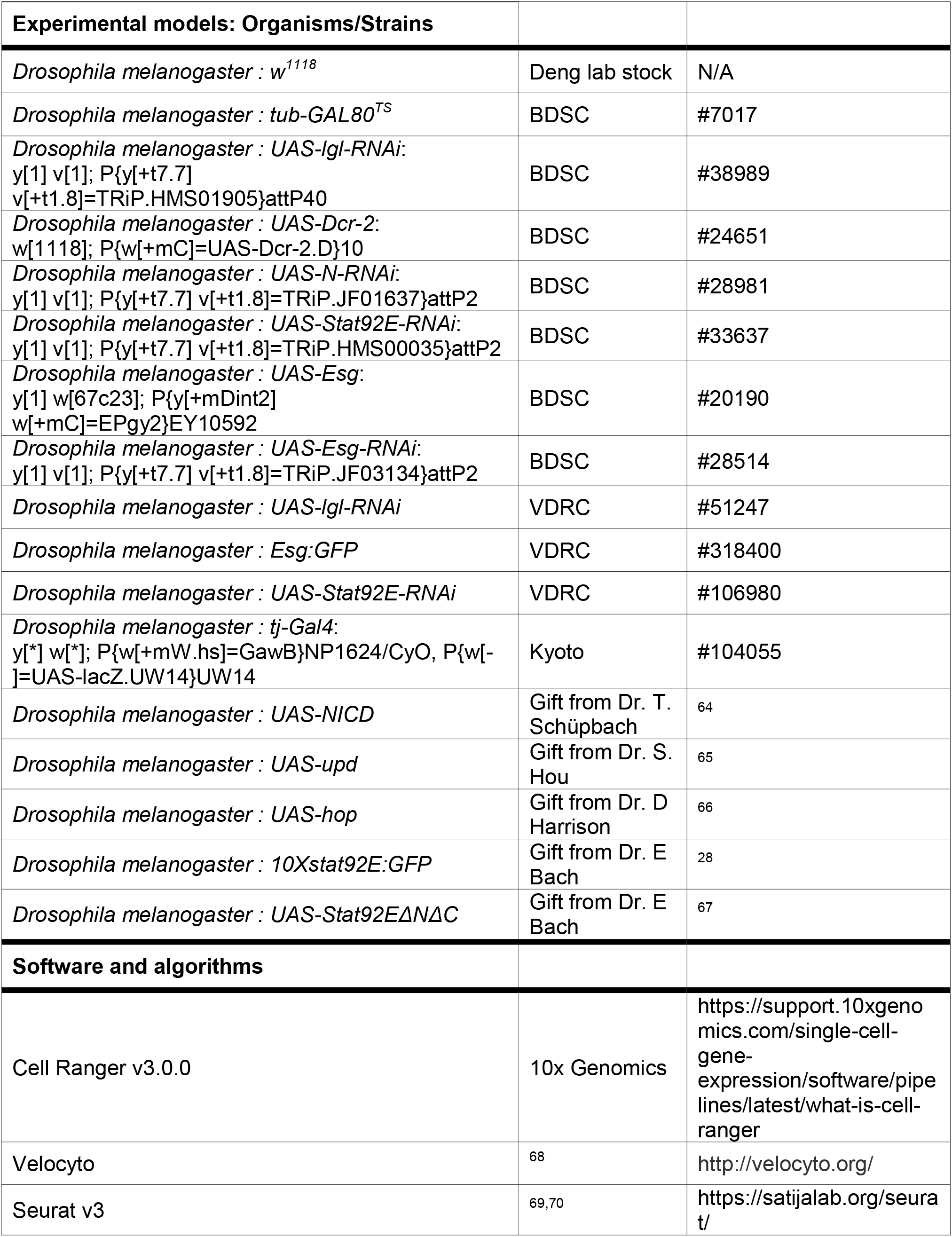
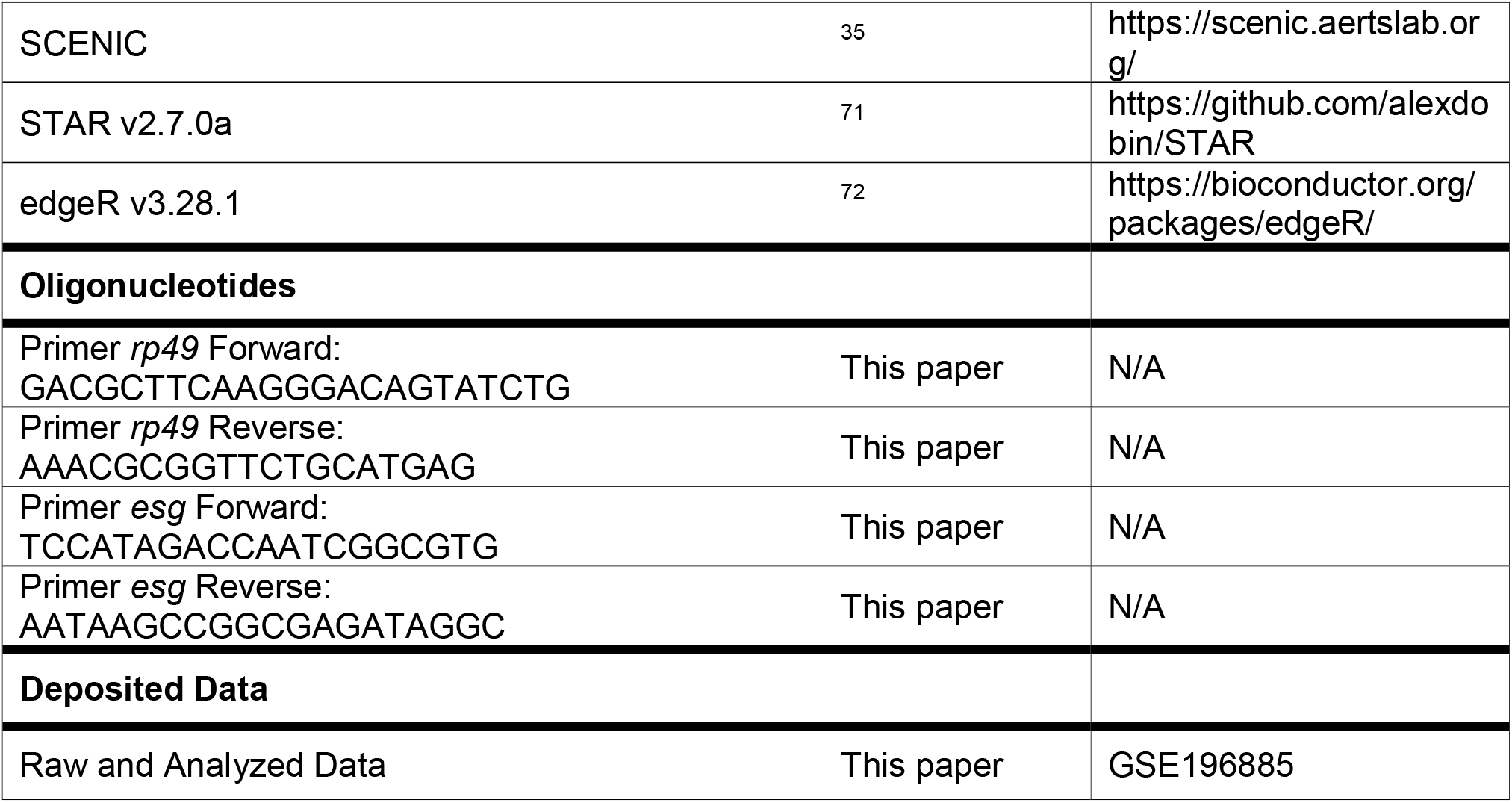

#### Resource availability

##### Lead contact

Further information and requests for resources and reagents should be directed to and will be fulfilled by the Lead Contact, Wu-Min Deng.

##### Materials availability

*Drosophila* lines described in this study are available from the Bloomington *Drosophila* Stock Center (BDSC), the Vienna *Drosophila* Resource Center (VDRC), or from the Lead Contact. Antibodies are available from the sources listed in the key resources table.

##### Data and code availability

No software or custom code was generated for this study. The whole-tissue and single-cell RNA-Seq datasets can be found in the GEO database under the identifier GSE196885.

##### Experimental model and subject details

Fly genetics and temperature shift information

*Drosophila* lines were maintained and crossed at 25°C on the BDSC cornmeal food (https://bdsc.indiana.edu/information/recipes/bloomfood.html). Flies were fed dry yeast 2 days before dissections to enhance oogenesis. For temperature-sensitive experiments, flies were maintained at 18°C and transferred to 29°C for transgene induction. For FLPout experiments, flies were reared at 25°C and were heat-shocked for 20 minutes at 37°C.

##### Immunohistochemistry, imaging and quantitative measurements

Ovaries were dissected in phosphate buffer saline (PBS) and fixed in 4% paraformaldehyde for 15 minutes. After washing with PBS and 0.2% Triton X-100 (PBT) and blocking with PBT and 0.5% BSA and normal goat serum, samples were incubated overnight with primary antibodies at 4°C with shaking. Antibody-stained tissues were washed 3 times with PBT again the next day and were subsequently incubated with secondary antibodies diluted in blocking solution for 2h at room temperature. Nuclei were labeled with DAPI (1:1000) and F-actin were labeled with Phalloidin (1:50). Samples were then mounted and imaged with the Zeiss LSM 800 Confocal Microscope were processed for figures using FIJI-ImageJ. Fiji was used for image analyses and processing. DAPI (1:20000) was used for nuclear DNA staining, which was used to identify nuclear area for each cell. Nuclear area and fluorescence intensity for specific channels were then measured quantitatively by the function “integrated density” in Fiji.

##### Fluorescence-activated cell-sorting (FACS) analysis

Analysis of a cell’s DNA content using flow cytometry has been described previously ^73^. Cell-dissociation protocol and library preparation were also performed as previously described (Jevitt, Chatterjee et al., 2020). Samples were treated with 50 U/ml papain with intermittent disruption via pipetting at room temperature until tissues were fully dissociated. Cell suspensions were passed through a 40μm filter and pelleted at 3,500 rpm for 12 minutes in an Eppendorf MiniSpin, following which, the samples were fixed with 4% paraformaldehyde with Vybrant DyeCycle Violet Stain (1:500, Life Technologies) for 30 minutes and washed twice with EBSS. Cell ploidy was determined by a flow cytometer (FACSAria, Becton Dickson) based on the excitations of Vybrant DyeCycle stain at 407 nm and GFP at 488 nm. CS&T beads (BD Biosciences) and SPHERO Rainbow Fluorescent Particles (Spherotech) were used as the calibration standards.

Whole-tissue samples were used for FACS analysis, since multilayers couldn’t be sorted using specific marker expressions. All quantifications associated with flow cytometry specifically focus on the cells that pass the gating thresholds (Q2) as shown in Supplementary Figure S2.

##### Quantitative Real Time (qRT)-PCR

Whole ovaries were lysed in TRIzol (Invitrogen) and RNAs were prepared according to the manufacturer’s protocol and quantified using nanodrop. For each sample, that were prepared in triplicates, 1μg of mRNA was reverse-transcribed using oligo-dT-VN primers and ImProm-II (Promega) as the reverse transcriptase. Real-time quantitative amplification of RNA was performed using the Sybr Green qPCR Super Mix (Invitrogen) and the iQ™5 Real-Time PCR Detection System from BioRad according to the manufacturer’s protocol. The relative expression of indicated transcripts was quantified with the CFX_Manager Software (BioRad) using the 2[-ΔΔC(T)] method. According to this method, the C(T) values for the expression of each transcript in each sample were normalized to the C(T) values of the control mRNA (*rp49*) in the same sample. The values of untreated cell samples were then set to 100% and the percentage transcript expression was calculated.

##### Whole-tissue RNA-Sequencing sample preparation

Total RNAs were extracted from ovaries after 24h and 96h of transgene expression in follicle cells. NEBNext Poly(A) mRNA Magnetic Isolation Module (NEB#E7490) and NEBNext Ultra II Directional RNA library Prep Kit for Illumina were used for library preparation. Libraries were sequenced using Rapid Run OBCG single read 50 bp on the Illumina Hi-Seq 2500 system. Following quality control (FastQC), raw reads were mapped to the *Drosophila melanogaster* Release-6 (BDGP6.28.dna.toplevel) reference genome (STAR). Counts of each gene were generated using featureCounts and the downstream analyses were performed using PCAtools and edgeR packages.

##### Single-cell RNA Sequencing

The cell-dissociation protocol and library preparation were performed as previously described (Jevitt, Chatterjee et al., 2020). Raw sequences from each 10X Genomics Chromium single-cell 3’ RNA-seq libraries were processed using Cell Ranger, that was followed by Velocyto to include counts of both spliced and unspliced reads. Seurat was then used for the processing of the output from Velocyto. Each sample was filtered for low-quality cells by setting sample-specific thresholds for UMIs, counts and mitochondrial gene-expression. For the *w^1118^* dataset, cells containing less than 13,000 UMIs, 2,000 genes and 8% mitochondrial gene percentage per cell but more than 650 genes were removed. Similarly, cells of the Lgl-KD dataset containing a maximum of 15,000 UMIs, 1,800 genes and 4.5% mitochondrial percentage but at least 750 genes per cell, and those of the Lgl-KD+Upd-OE dataset containing less than 8000 UMIs, 1,250 genes and 5% mitochondrial percentage but more than 2,500 UMIs and 700 genes per cell were also removed. As a result, 19,986 cells were obtained for the *w^1118^* control, 16,060 cells for Lgl-KD and 42,235 cells for Lgl-KD+Upd-OE datasets for downstream processing. For the integration of follicle cells from separate datasets, cells are aligned using 2000 anchor genes. Following UMAP embedding (using 60 Principal Components; resolution=1), non-epithelial clusters were removed, along with clusters that constitute less than 0.2% of the total cell number. The final dataset was curated iteratively. ALRA imputation of the sparse counts for genes in the “spliced” assay was applied to recover missing values using the suggested default settings, and the imputed counts were only used for visualization. Regulon activity was assessed using standard SCENIC pipeline and *Drosophila-specific* database for motif rankings (cisTarget v8 motif collection dm6-5kb-upstream-full-tx-11species.mc8nr).

##### Statistics and reproducibility

Statistical analyses and graphs were prepared in R. Unpaired t-tests was used for 2-sample comparisons. Results were confirmed after at least 2 independent experiments and each experiment involved the dissection of at least 25 flies (25 pairs of ovaries) per sample. Two replicates were used for each whole-tissue RNA-Seq dataset, the count matrix from which were then compared with the aggregated counts of unfiltered single-cell RNA-Seq datasets using PCA analysis (Supplementary Figure S3).

Samples for whole-tissue RNA-Seq were subjected to 24h and 96h of transgene expression, while those for single-cell RNA-Seq were subjected to 72h to promote the relative abundance of detectable phenotypes in the whole-tissue RNA-Seq and to avoid excessive death in the single-cell RNA-Seq experiments, respectively.

## EXCEL TABLE TITLES AND LEGENDS

**Supplemental Table 1:** Differential gene expression analyses using whole-tissue RNA-Seq datasets. Related to Figures 2–3.

**Supplemental Table 2:** Enriched motifs over-represented in the *tj^TS^>lgl^RNAi^+Upd* (72h) dataset. Related to Figure 5.

**Supplemental Table 3:** Quantifications of the qRT-PCR experiment to detect Esg mRNA. Related to Figure 6.

---

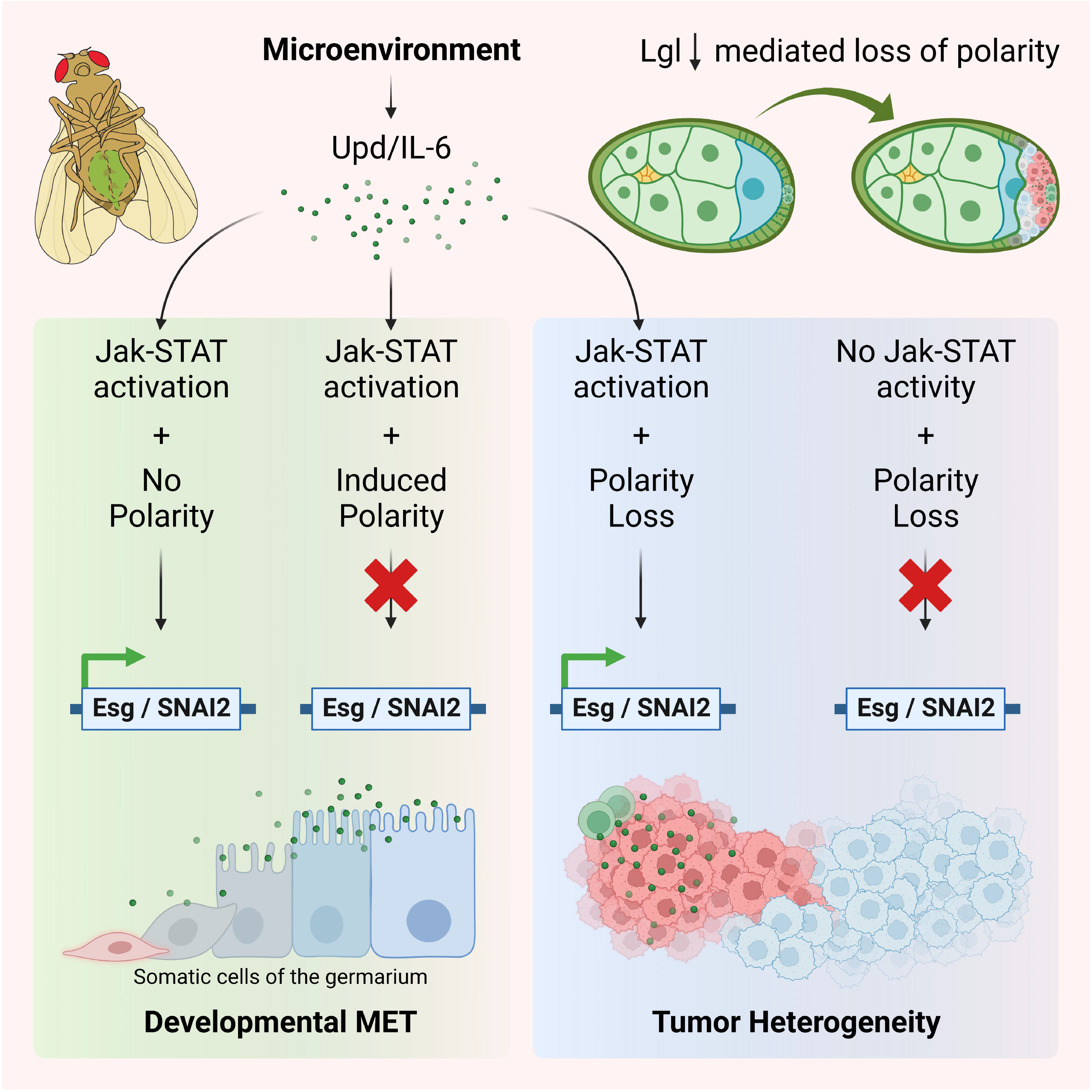

