## Supplementary Figures for "Cell polarity opposes Jak-STAT mediated Escargot activation that drives intratumor heterogeneity in a *Drosophila* tumor model"

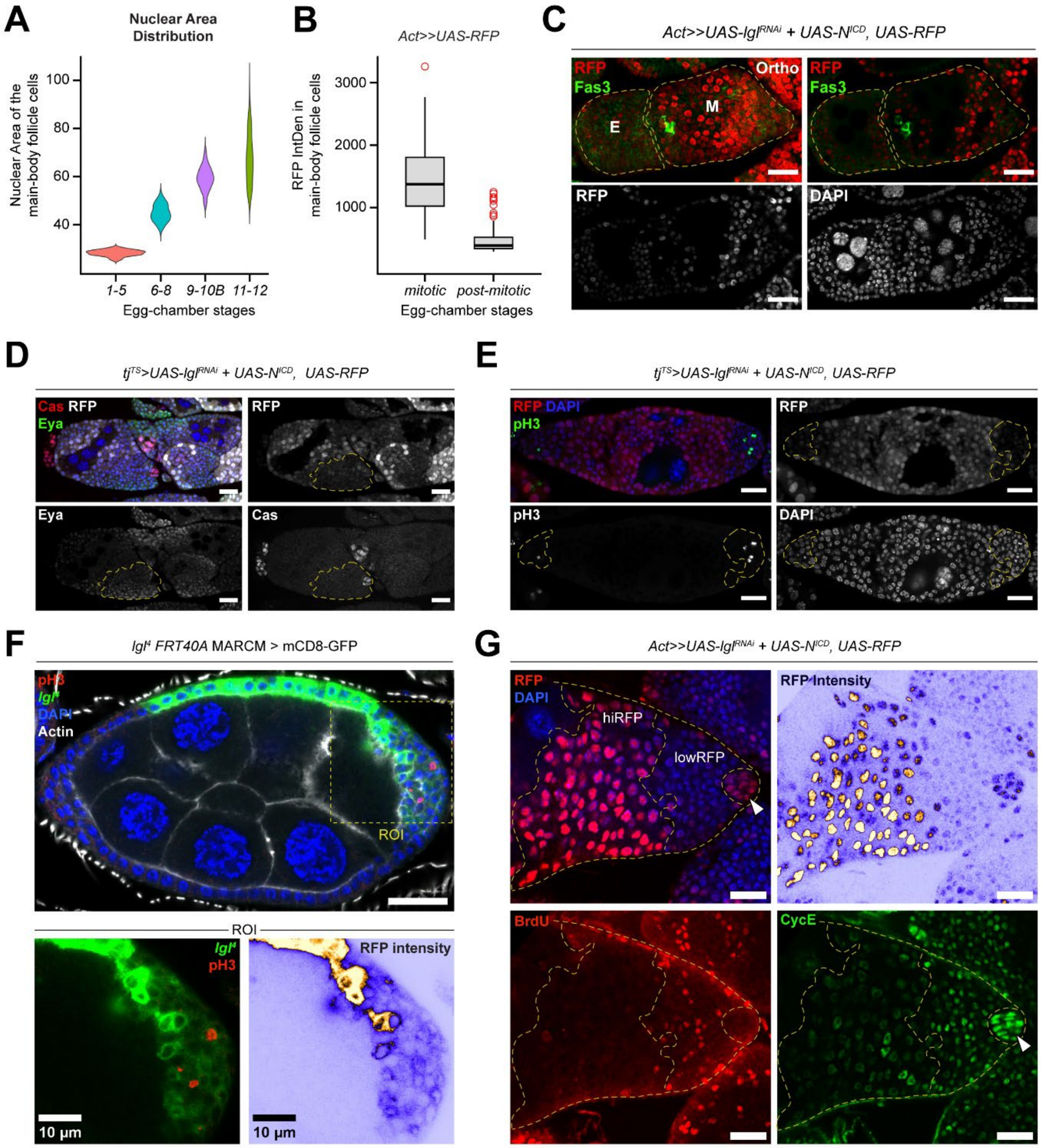


**Figure S1. RFP heterogeneity does not depend on the choice of genotype, driver or fluorescent marker and shows cell-cycle differences, Related to Figure 1. (A)** Violin plot to show the developmental-stage (X-axis) specific distribution of nuclear area of main-body follicle cells (Y-axis). **(B)** Box-and-whisker plot to show the distribution of RFP intensity (Y-axis) in mitotic and post-mitotic follicle cells (X-axis) from egg chambers driving RFP under the influence of the *Actin*-FLPout promoter. **(C)** Confocal images to show multilayers of RFP-expressing *Actin*-FLPout Lgl-KD+$N^{ICD}$ cells. Differential RFP expression (red) is observed between follicle cells at early- (E) and mid-stage (M) oogenesis. Polar cells are marked by high Fas3 expression (green). Ortho = Orthogonal projection. Discrete egg chambers are shown within dashed lines. **(D)** Confocal images to show the expression of Eya (green) and Cas (red) in multilayered Lgl-KD+$N^{ICD}$ cells. Expression of Eya, Cas and RFP are also shown separately. Posterior multilayer is identified within the dashed yellow line. **(E)** Confocal images to show the expression of the mitotic marker phosphor-Histone3 (pH3) (green) within the multilayered Lgl-KD+$N^{ICD}$ cells (identified within the dashed yellow line). Expression of RFP, DAPI and pH3 are also shown separately. **(F)** Confocal image to show the homozygous clones of $lgl^4$ (loss-of-function *lgl* mutant), identified by membrane-bound GFP expression (green). A Region of Interest (ROI), highlighted within a dashed yellow square, is magnified and shown separately, where discrepancy in RFP intensity is shown as a color spectrum where yellow represents high and blue represents low intensities. Mitotic division in the multilayered follicle cells is indicated by pH3 expression (red). **(G)** Confocal image to show relative enrichment of CycE and BrdU staining in the posterior multilayers of follicle cells. CycE expression is indicated in green and BrdU incorporation is indicated in red. RFP intensity is shown as the blue-yellow spectrum. $RFP^+$ (red) follicle cells are divided into high-RFP expressing (hiRFP) and low-RFP expressing (lowRFP) cells. G2-arrested polar cells (white arrowhead) are identified by CycE enrichment, morphology and location within the multilayer. Low-RFP cells exhibit a higher flux in cell-cycle phase as differential enrichment of CycE and increased BrdU incorporation is detected. Nucleus is marked by DAPI staining. CycE staining and BrdU incorporation was performed as described in Wang et al. (2020)[1] and Zielke et al. (2011)[2], respectively. Scale bars: 20μm, unless mentioned otherwise.

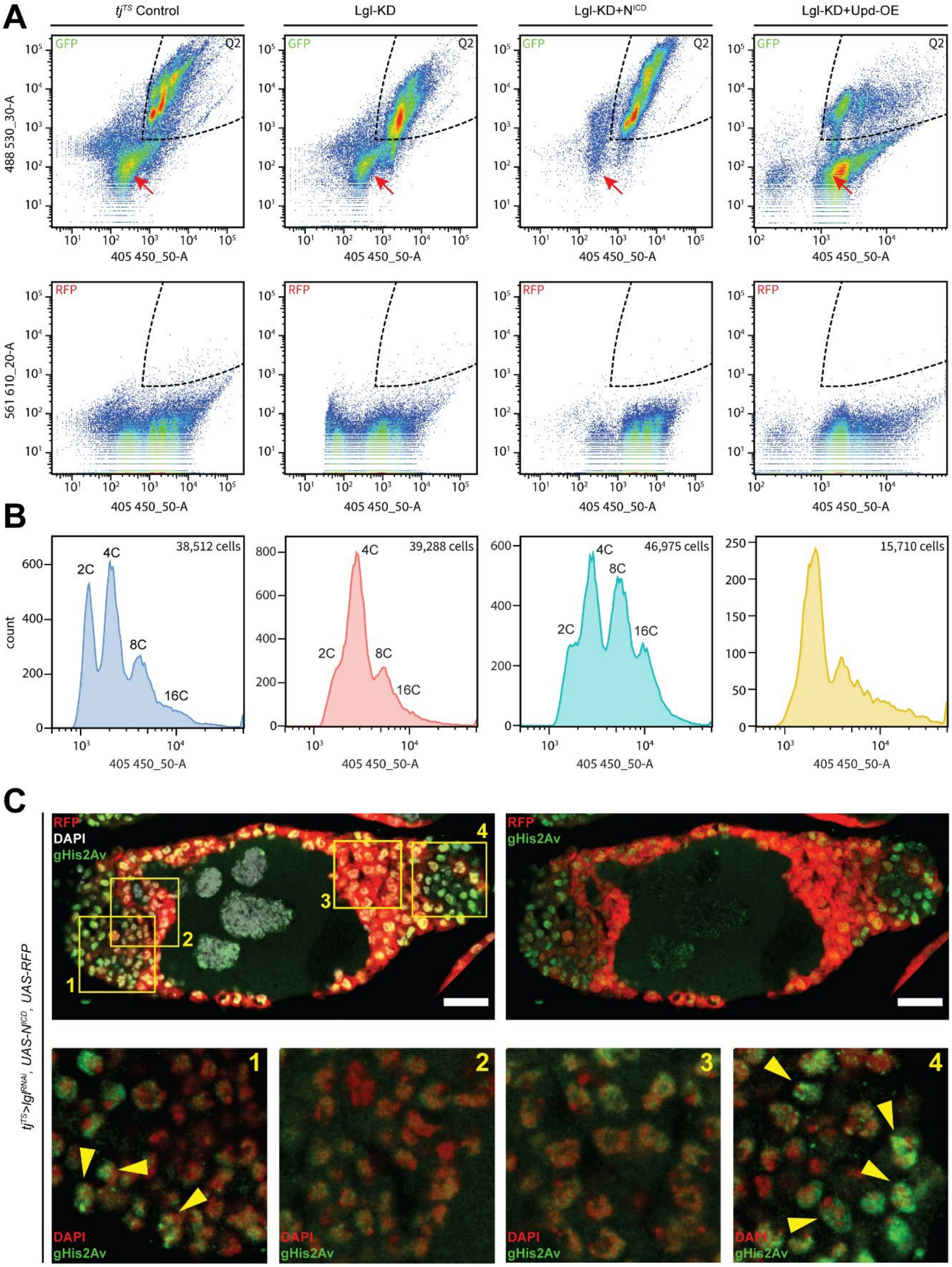
A
tj^TS Control
Lgl-KD
Lgl-KD+N^ICD
Lgl-KD+Upd-OE
GFP
Q2
488 530_30-A
405 450_50-A
RFP
561 610_20-A
B
38,512 cells
39,288 cells
46,975 cells
15,710 cells
2C
4C
8C
16C
count
C
RFP
DAPI
gHis2Av
tj^TS>lgl^RNAi, UAS-N^ICD, UAS-RFP
1
2
3
4

**Figure S2. FACS analysis of GFP$^+$ follicle cells from relevant genotypes driven by *tj*$^{TS}$>GAL4 driver, Related to Figure 1. (A)** Scatter plot of flow cytometry analysis to identify GFP$^+$ follicle cells in control, Lgl-KD, Lgl-KD+N$^{ICD}$ and Lgl-KD+Upd-OE egg chambers, where the gating strategy and distribution of GFP$^+$ cells (top) and background (bottom) is shown. Note that the gates were set up by dividing the distribution into quadrants and only selecting the quadrant (Q2) that contained GFP$^+$ cells but did not include background, as was identified by detecting RFP which was absent from the genotypes. Likely cluster of germline cells are indicated by the red arrow. **(B)** Distribution of ploidy in cells within Q2 for each genotype. Expected ploidy for each peak is indicated. **(C)** Confocal images to show the multilayered egg chamber containing Lgl-KD+N$^{ICD}$ follicle cells at midoogenesis. The low RFP expressing multilayered cells exhibit foci of γ-His2Av (or gHis2Av) staining (green), as marked by yellow arrowheads in the 4 subsetted Region of Interests (ROIs 1-4, indicated by yellow squares). ROIs 1 and 4 mostly consist of follicle cells having low intensities of RFP expression, while ROIs 2 and 3 mostly contain cells expressing RFP at high intensities. In contrast to the top panel where RFP expression is shown in red, the bottom panel shows DAPI staining in red to allow a uniform contrast for the assessment of γ-His2Av staining. Nucleus is marked by DAPI staining. Scale bars: 20μm.

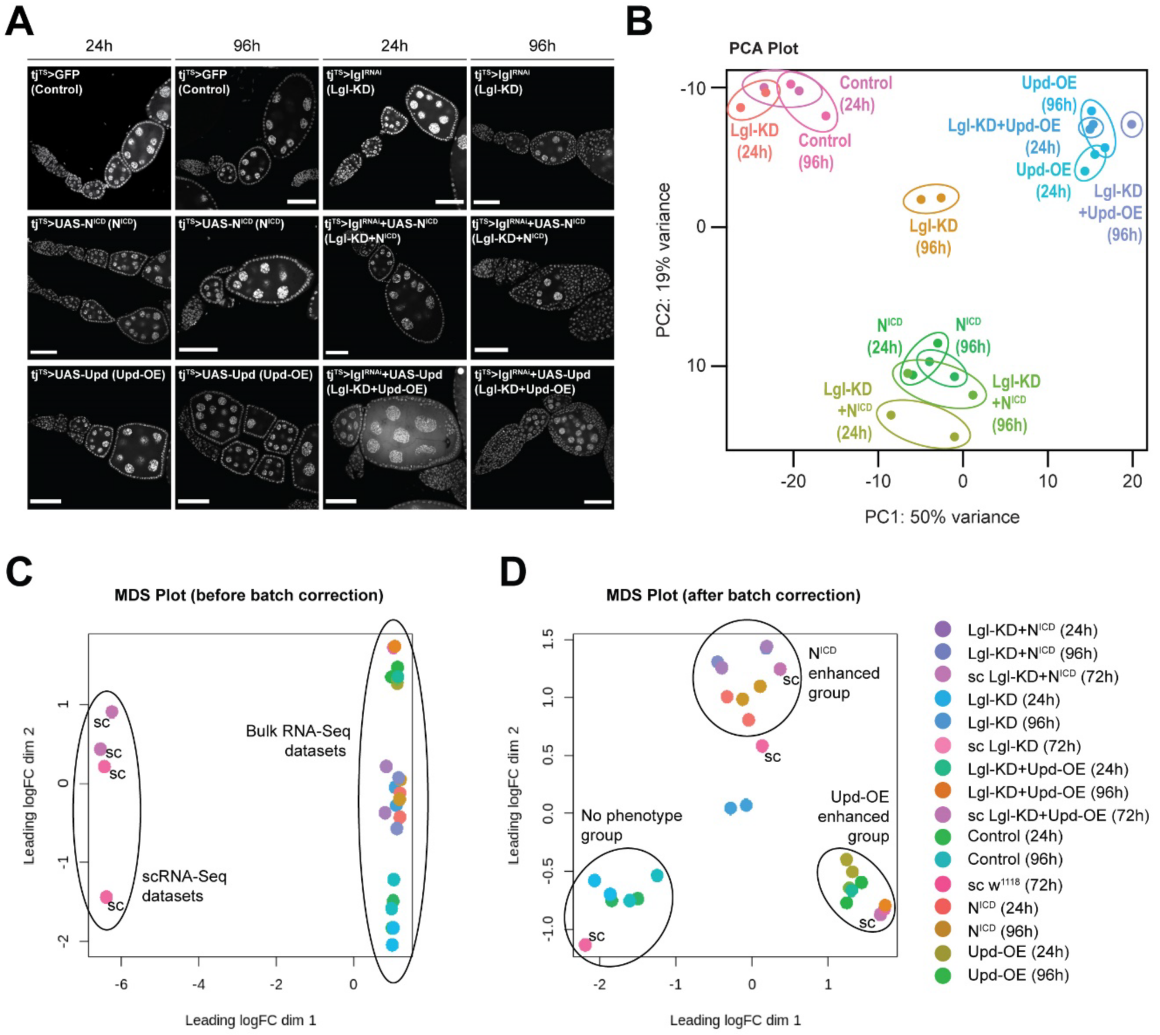


**Figure S3. Analysis of whole-tissue RNA-Seq datasets, Related to Figure 2 and 3. (A)** Confocal images of representative samples of ovarioles containing follicle cells of relevant genotypes driven by the *tj$^{TS}$*>GAL4 driver after 24h and 96h of transgene expression. These samples were used as inputs for whole-tissue RNA-Seq experiment. Nucleus is marked by DAPI staining. Scale bars: 20μm. **(B)** PCA plot to show the relative distribution of whole-tissue RNA-Seq samples. Each sample consists of 2 replicates which are color-coded and contained within the ellipses. The sample names are mentioned next to the corresponding ellipses. **(C)** Multi-Dimensional Scaling (MDS) plot depicting the relative distribution of bulk and single-cell RNA-Seq samples (marked by "sc" next to the representative dots) on reduced dimensions. Due to the inherent differences in time-points, sequencing depth, cell types and undefined technical differences, the samples are largely separated by batch effects. Note that the singe-cell datasets only contain follicle cells, as the other positively-identified cell types have been removed from the analysis. **(D)** MDS plot of the same samples after batch correction shows co-clustering of samples belonging to the same genotype, irrespective of technical differences. This co-clustering adds validation to the fidelity of transcriptomic datasets and their similarities across time-points and within genotypes.

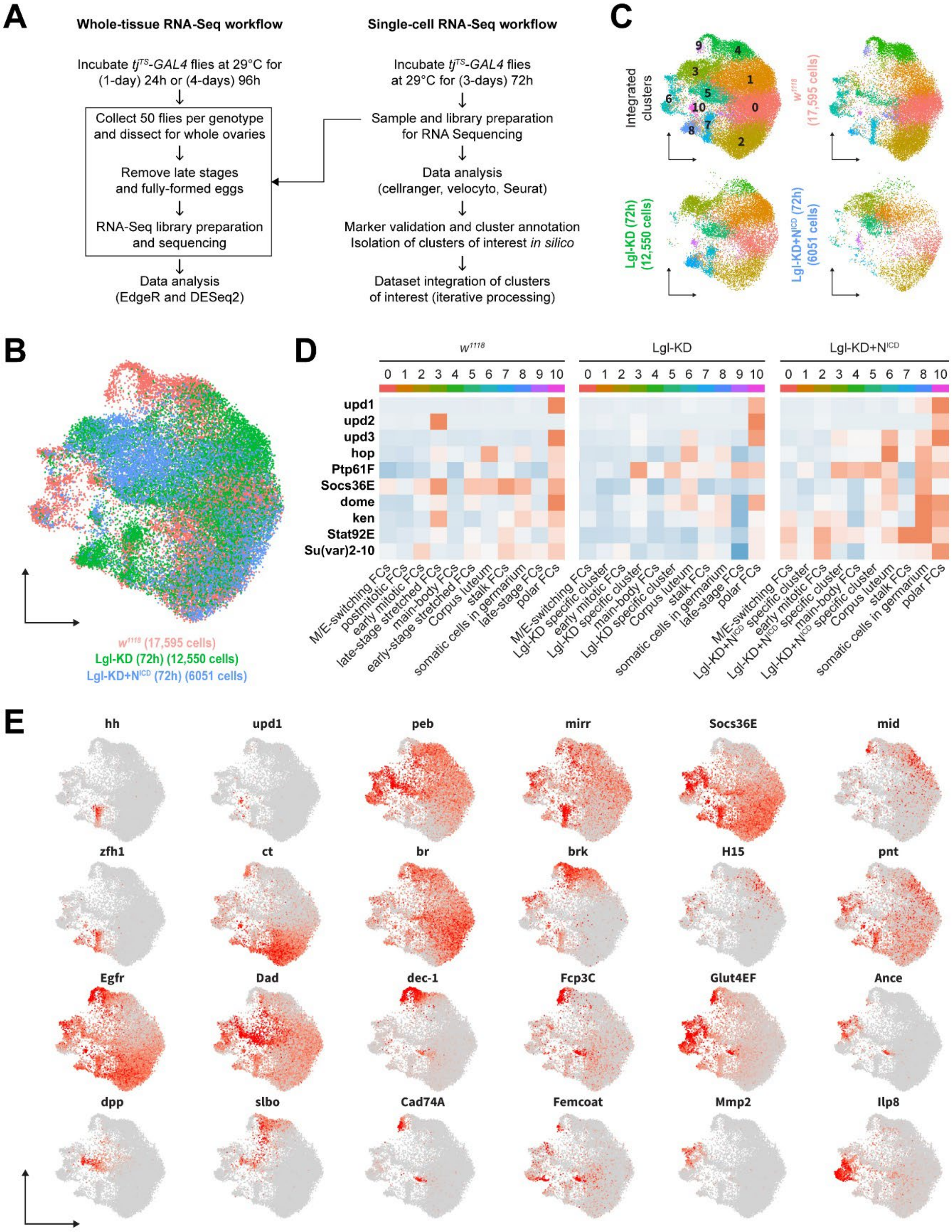
A
Whole-tissue RNA-Seq workflow
Incubate tjTS-GAL4 flies at 29°C for (1-day) 24h or (4-days) 96h
Collect 50 flies per genotype and dissect for whole ovaries
Remove late stages and fully-formed eggs
RNA-Seq library preparation and sequencing
Data analysis (EdgeR and DESeq2)
Single-cell RNA-Seq workflow
Incubate tjTS-GAL4 flies at 29°C for (3-days) 72h
Sample and library preparation for RNA Sequencing
Data analysis (cellranger, velocyto, Seurat)
Marker validation and cluster annotation
Isolation of clusters of interest in silico
Dataset integration of clusters of interest (iterative processing)
B
w1118 (17,595 cells)
Lgl-KD (72h) (12,550 cells)
Lgl-KD+NICD (72h) (6051 cells)
C
Integrated clusters
w1118 (17,595 cells)
Lgl-KD (72h) (12,550 cells)
Lgl-KD+NICD (72h) (6051 cells)
D
w1118
Lgl-KD
Lgl-KD+NICD
upd1
upd2
upd3
hop
Ptp61F
Socs36E
dome
ken
Stat92E
Su(var)2-10
M/E-switching FCs
postmitotic FCs
early mitotic FCs
late-stage stretched FCs
main-body FCs
early-stage stretched FCs
Corpus luteum
stalk FCs
somatic cells in germarium
late-stage FCs
polar FCs
Lgl-KD specific cluster
Lgl-KD+NICD specific cluster
E
hh
upd1
peb
mirr
Socs36E
mid
zfh1
ct
br
brk
H15
pnt
Egfr
Dad
dec-1
Fcp3C
Glut4EF
Ance
dpp
slbo
Cad74A
Femcoat
Mmp2
Ilp8

**Figure S4. Analytical pipeline for whole-tissue and single-cell sequencing and scRNA-Seq dataset integration, Related to Figure 2. (A)** Schematics of whole-tissue and single-cell RNA-Seq pipelines employed in this study. **(B)** UMAP plot to show the integrated embedding of $w^{1118}$ follicle cells (red) with those expressing Lgl-KD (green) and Lgl-KD+$N^{ICD}$ (blue). **(C)** UMAP plot to show the embedding of follicle cells, split by genotype (as indicated). Cells are colored by their cluster identities (identified within the integrated embedding in the top-left panel). **(D)** Heatmap to show the average expression (in cells per cluster) of genes belonging to the Jak-STAT signaling pathway in the individual datasets[3]. The expression of genes is scaled between +2 (red) and -2 (blue) log2FC. **(E)** Gene enrichment plots to show the expression of known markers used to annotate cell-type identities of clusters in the integrated dataset.

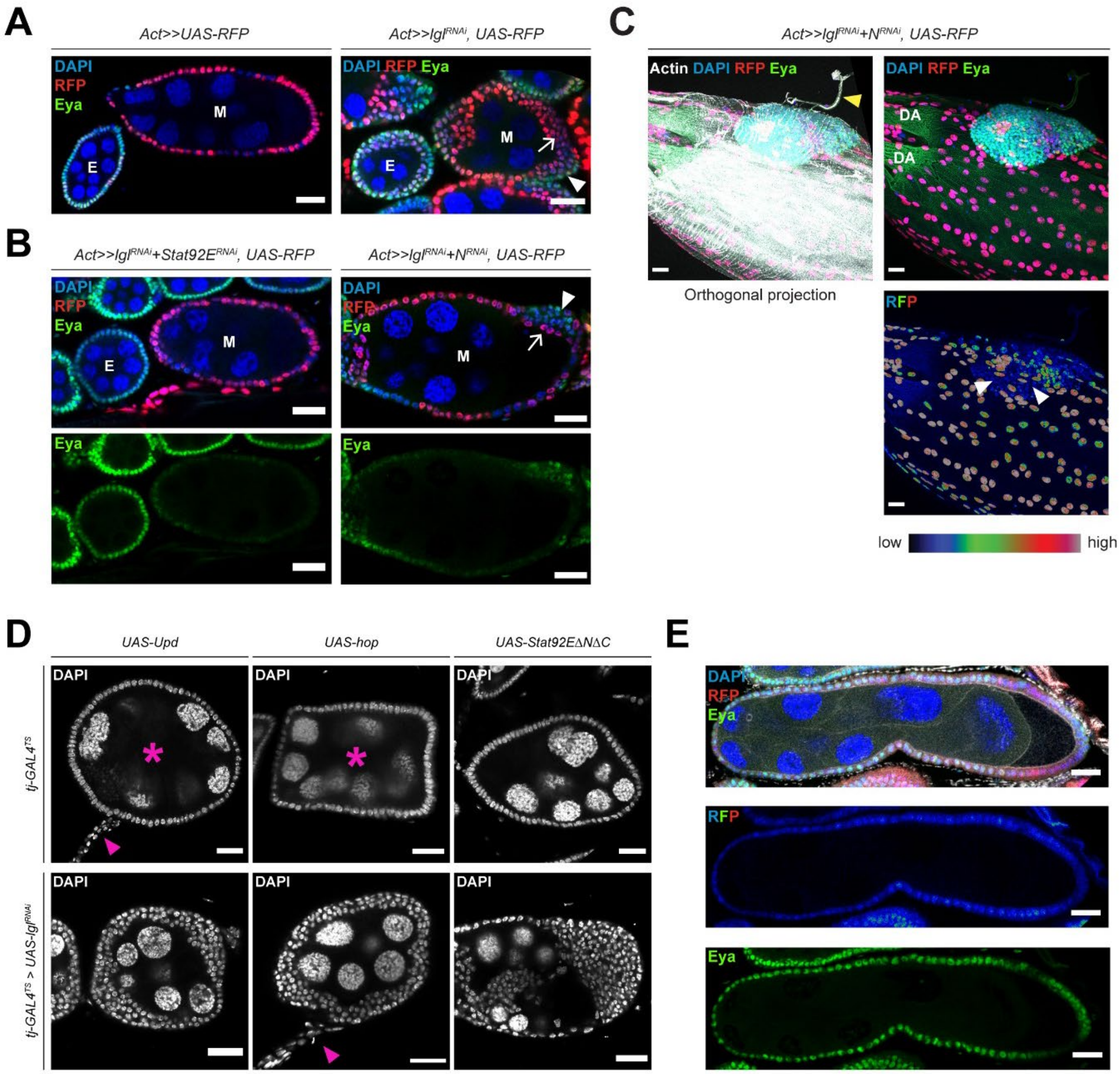


**Figure S5. Jak-STAT signaling affects immature cells in the multilayer, Related to Figure 3. (A)** Confocal images showing the expression of Eya (green) and RFP (red) discrepancy in the multilayers. In the left panel, egg chambers at Early (E) oogenesis contain immature Eya$^+$ follicle cells (low RFP), while mature follicle cells in egg chambers at midoogenesis (M) do not express Eya and show high RFP expression. The right panel shows multilayered Lgl-KD follicle cells in egg chambers at both E and M-stages. Note that while all the follicle cells at early oogenesis exhibit Eya expression, the multilayered cells contain both Eya$^+$ and Eya$^-$ cells. The Eya$^-$ cells (marked by arrow) show higher RFP intensity, compared to the Eya$^+$ cells (marked by arrowhead). **(B)** Confocal images of egg chambers with Stat92E-KD (left) and N-KD (right) in RFP$^+$ Lgl-KD follicle cells. Multilayers are completely rescued in egg chambers containing Lgl-KD+Stat92E-KD follicle cells. Eya expression is observed in egg chambers at E, but not at M-stages, except in egg chambers containing Lgl-KD+N-KD follicle cells, where multilayers of RFP$^+$ cells are rescued (marked by arrow) while Eya$^+$ cells persist basally in the multilayers (marked by arrowhead). **(C)** Confocal images showing the orthogonal projection of a stage-12 egg chamber) containing RFP$^+$ (red) Lgl-KD+N-KD follicle cells (18.5% egg chambers with Lgl-KD+N-KD follicle cells formed intact late-stage egg chambers (n=23/124). The top panels show the presence of Eya$^+$ (green) cells on the basal side of the epithelial layer of the egg chamber. F-Actin is stained by Phalloidin (white), and a trachea-like outgrowth from the Eya$^+$ cells is indicated by a yellow arrowhead. Dorsal Appendages (DA) are indicated on the image. Bottom panel highlights the discrepancy in RFP intensity, where Eya$^+$ multilayered cells (marked by white arrowheads) show decreased intensities of RFP (which is shown as a spectrum as indicated below). **(D)** Confocal images of egg chambers at midoogenesis upon Upd-OE, Hop-OE and Stat92EΔNΔC expression in follicle cells with (upper panel) and without Lgl-KD (lower panel). Multilayering is only detected in egg chambers with Lgl-KD follicle cells. Abnormal egg-chamber shape is indicated by magenta asterisks (*), while abnormally-long stalk cells are indicated by magenta arrowheads. **(E)** Confocal images of egg-chambers at Stage 8 (as estimated by measuring the germline-nuclei volume) showing Eya$^+$ (green) follicle cells, that also show low RFP intensity. Nucleus is marked by DAPI staining. Scale bars: 20μm.

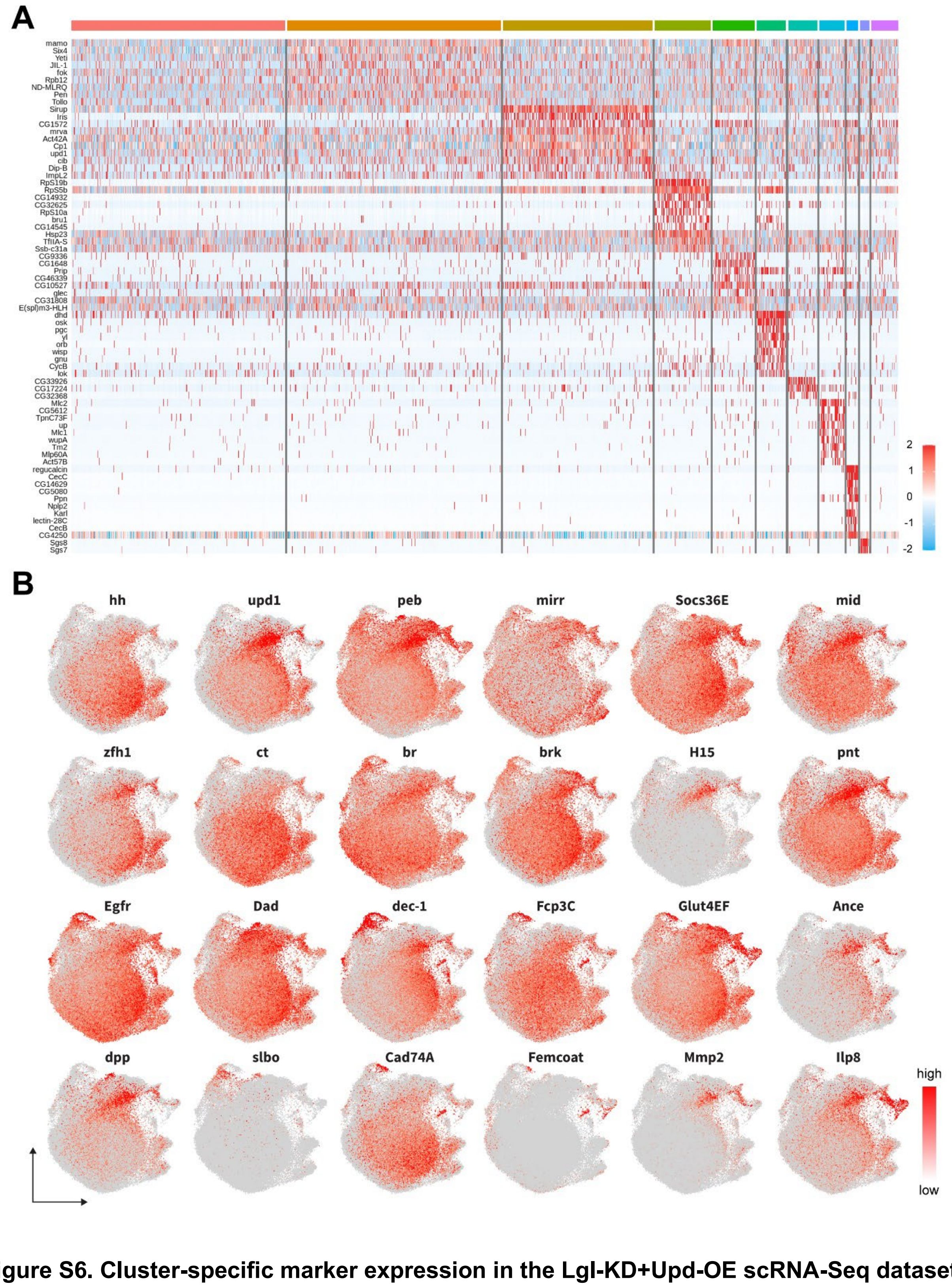


**Figure S6. Cluster-specific marker expression in the Lgl-KD+Upd-OE scRNA-Seq dataset and expression of known markers in the Integrated dataset, Related to Figures 4 and 5.**

**(A)** Heatmap to show the cluster-specific marker expression in the Lgl-KD+Upd-OE scRNA-Seq dataset. The expression of genes is scaled between +2 (red) and -2 (blue) log2FC. **(B)** Gene enrichment plots showing the cluster-specific enrichment of known markers used to annotate cell-type identities of each cluster in the integrated dataset of $w^{1118}$, Lgl-KD, Lgl-KD-$N^{ICD}$ and Lgl-KD+Upd-OE samples.

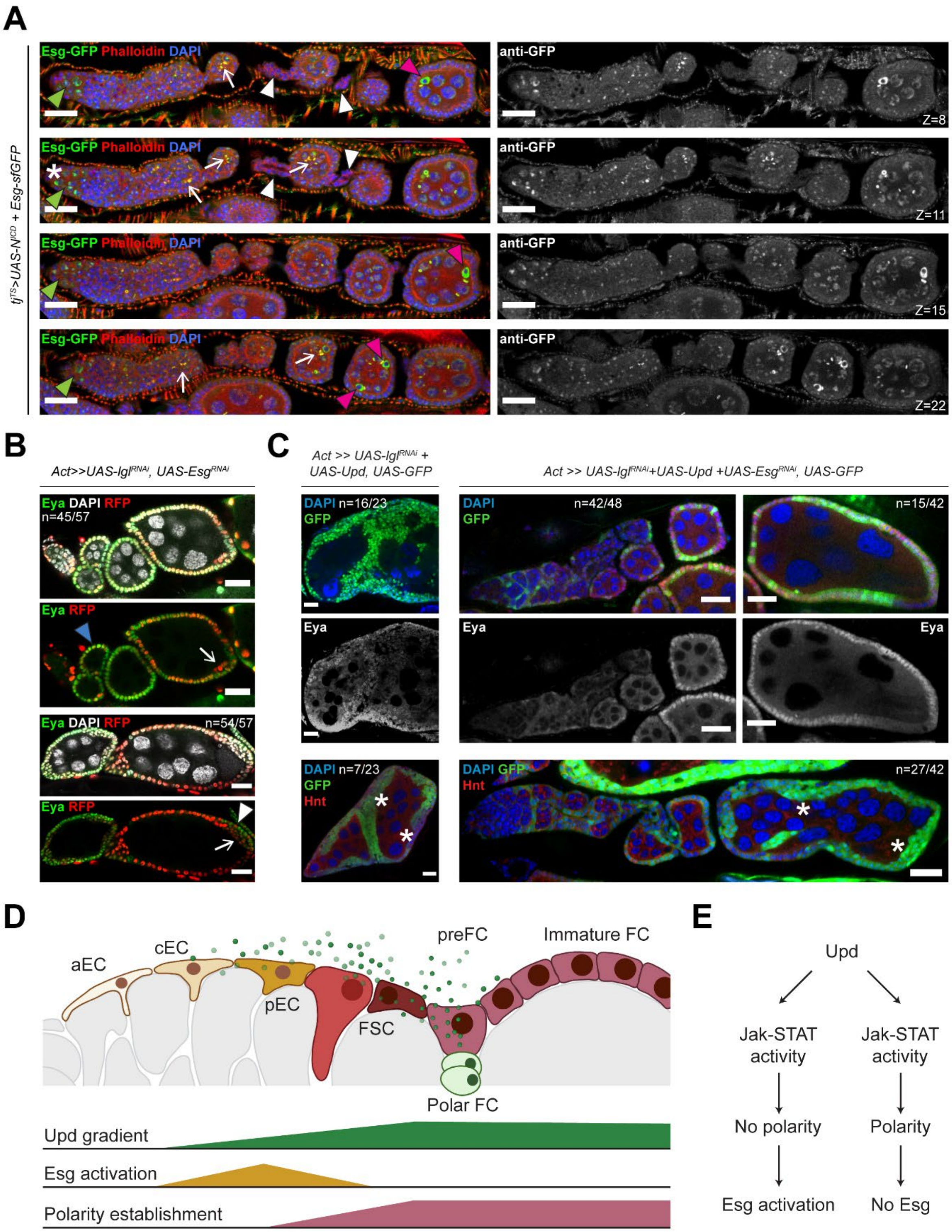


**Figure S7. Esg expression in ovarioles containing Lgl-KD+$N^{ICD}$ follicle cells and the**

**impact of Esg-KD in multilayered and normal egg chambers, Related to Figure 7. (A)** Confocal images to show the localization of Esg-sfGFP in egg chambers containing Lgl-KD+$N^{ICD}$ follicle cells during early oogenesis at different Z planes (indicated at the bottom-right corner of images in the right panel). Green arrowhead marks the $GFP^{+}$ germline cyst cells within the germarium, while the asterisk marks the absence of GFP in the germline stem cells. The arrows mark the ring canals, where GFP expression (green) coincide with high F-Actin (red). Abnormal stalk cells show no localization of Esg (indicated by the absence of GFP) and are marked by white arrowheads. Germline-cell encapsulation error is shown in egg chambers containing multiple oocytes, as marked by the magenta arrowheads. This staining pattern is similar to that observed in the control egg chambers with no morphological defects. Nuclei are stained with DAPI (blue). **(B)** Confocal image to show partial rescue of Lgl-KD multilayering by Esg-KD in $RFP^{+}$ follicle cells (red). Eya staining is shown in green, while nuclei are stained with DAPI (white). Complete rescue of multilayering is observed in early-stage egg chambers (identified by blue arrowhead). Mature delaminated cells are identified by arrows, while monolayered immature cells are indicated by white arrowhead. **(C)** Confocal image to show severely-disrupted FE of egg chambers with GFP+ (green) Lgl-KD+Upd-OE follicle cells alone (Left) and those co-expressing Esg-KD (Right). Middle row shows Eya staining (grayscale) to mark immature follicle cells. Oocytes within compound egg chambers are indicated by asterisks (*). Nuclei are stained with DAPI (blue). Scale bars: 20μm. **(D)** Inferred distribution of Upd-mediated Jak-STAT activation, cell-polarity establishment and Esg activation within the somatic cells of the germarium that represent the earliest stages of follicle-cell lineage. **(E)** Model to show the likely relationship between cell polarity, Jak-STAT activation and Esg activation in the somatic cells within the germarium.
